# Metal microdrive and head cap system for silicon probe recovery in freely moving rodent

**DOI:** 10.1101/2020.12.20.423655

**Authors:** Mihály Vöröslakos, Peter C. Petersen, Balázs Vöröslakos, György Buzsáki

## Abstract

High-yield electrophysiological extracellular recording in freely moving rodents provides a unique window into the temporal dynamics of neural circuits. Recording from unrestrained animals is critical to investigate brain activity during natural behaviors. The use and implantation of high-channel-count silicon probes represent the largest cost and experimental complexity associated with such recordings making a recoverable and reusable system desirable. To address this, we have designed and tested a novel 3D printed head-gear system for freely moving mice and rats. The system consists of a recoverable microdrive printed in stainless steel and a plastic head cap system, allowing researchers to reuse the silicon probes with ease, decreasing the effective cost, and the experimental effort and complexity. The cap designs are modular and provide structural protection and electrical shielding to the implanted hardware and electronics. We provide detailed procedural instructions allowing researchers to adapt and flexibly modify the head-gear system.

## Introduction

Action potentials are the common currency of communication between neurons and they can be detected as voltage fluctuation in the extracellular space (Adrian and Moruzzi, 1939). However, recording from representative ensembles of neurons simultaneously requires electrodes with multiple recording sites. Multi-wire twisted electrodes (tetrodes) and silicon probes offer the possibility to record tens to hundreds of neurons simultaneously from multiple cortical and subcortical structures simultaneously in freely moving animals (Blanche et al., 2005; Buzsáki, 2004; Csicsvari et al., 2003; Jun et al., 2017; McNaughton et al., 1983; Montgomery et al., 2008; Wise and Najafi, 1991). For cost benefits, microwire arrays are a popular choice for neuroscientists, despite the amount of manual labor involved (Edell et al., 1992). Silicon probes, while more expensive, do not require assembly, the tissue-volume displacement is minimal (Buzsáki, 2004; Kipke et al., 2008), recording properties are consistent (site impedance and locations) and geometric configurations (number of shanks, distance, and pattern of recording sites) can be customized to suit the architecture of the particular brain structure under study (Scholvin et al., 2016; Wise and Najafi, 1991). The availability of high-channel-count electrophysiology amplifier chips (e.g., RHD-2132 and RHD-2164, Intan Technologies, Los Angeles, CA (Berényi et al., 2013)) and integrated designs (Jun et al., 2017) have accelerated the spread of large-scale recordings. μLEDs and micro fluidic delivery can also be integrated into silicon-based electrodes and can offer unique spatiotemporal control of neuronal activity (Wu et al., 2015; Kim et al., 2020).

Despite the availability of cutting-edge recording electrodes, the development of implantation techniques such as the Flexdrive, Shuttledrive, DMCdrive and the Hyperdrive (Voigts et al., 2013, 2020; Kim et al., 2020; Lu et al., 2018) has lagged behind. Electrodes are either fixed in brain tissue or attached to a microdrive to allow the advancement of the electrode after implantation (Chung et al., 2017, 2017; Fee and Leonardo, 2001; Korshunov, 2006; Vandecasteele et al., 2012; Wilson and McNaughton, 1993; Yamamoto and Wilson, 2008). Microdrives and accompanying head gear protection and shielding inevitably add extra weight (weight = 0.12 - 1 g, drives designed for mice) and volume (skull surface area = 7.68 – 252 mm^2^, drives designed for mice) to the implant (Table 1). The weight, volume and footprint of the microdrive can limit comfortable movement of small rodents and can prevent flexible multiregional recordings in mice (Headley et al., 2015). Yet, chronic recordings from freely behaving subjects are essential in many experiments, where the relationship between neuronal activity and movement, perception, learning and memory, decision making, and other forms of cognition are studied to disambiguate overt behavior and hidden variables (Juavinett et al., 2019; Jun et al., 2017; Steinmetz et al., 2020).

**Table 1.** Summary of microdrive designs used in mice.

| Study/Company | Width (mm) | Length (mm) | Height (mm) | Footprint (mm<sup>2</sup>) | Weight (g) | Travel distance (mm) | Easy recovery |
| --- | --- | --- | --- | --- | --- | --- | --- |
| Vandecasteele et. al. 2012 | 4.3 | 6.4 | 13 | 27.52 | 0.6 | 8-12 | no |
| Korshunov 2006 | 3.2 | 2.4 | 9.6 | 7.68 | 0.12 | 5 | no |
| Janelia Research Campus | 3.5 | 3.8 | 9 | 13.3 | 0.8 | 5 | no |
| Janelia Research Campus | 2.5 | 3.8 | 10 | 9.5 | 0.5 | 5 | no |
| Cambridge Neurotech | 2.5 | 4 | 9 | 10 | 0.54 | 5 | no |
| Neuronexus | 12.5 | 11.5 | 8.5 | 143.75 | 0.36 | 1 | no |
| NeuroNex MINT | 3.2 | 7.5 | 16 | 24 | 0.67 | 4.8 | yes |
| Chung et. al. 2017 | 6.26 | 5.26 | 9 | 32.93 | 0.4 | 2 | yes |
| Juavinett et. al. 2019 | 18 | 14 | 20 | 252 | 1 | - | yes |
| Voroslakos et. al. | 3.1 | 5 | 15.3 | 15.5 | 0.87 | 7 | yes |

An ideal microdrive should have movement precision, mechanical stability, minimal size, low weight, and the ability for flexible customization. Commercially available microdrives are expensive and hard to customize. Disposable 3-D printed customized drives and head gear have reduced costs (Allen et al., 2020; Chung et al., 2017; Headley et al., 2015). However, the most expensive component of the recording system is the silicon probe. Given the cost (from $1000 for 32-channel passive recording probes to > $3000 for μLED probes or larger channel count probes), the option of reusing silicon probes is, therefore, an important current goal (Juavinett et al., 2019). In addition to reducing costs, repeated use of the same probe/headgear would allow better consistency for recordings across animals, enhance data reproducibility, and reduce electrode/headgear preparation for surgery. Achieving this goal requires an integrated design of a reusable microdrive and head gear to increase recording stability and protect/shield sensitive drive and electronic components (Chung et al., 2017; Senzai et al., 2019). We report here the design and testing of an integrated 3D printed headgear system (including microdrives and protective head cap) for both mice and rats. Our design reduces surgery time substantially. The fast and reliable recovery of the probe and reuse of the same system in multiple animals decreases costs and experimenter effort.

## RESULTS

### Recoverable metal microdrive

3D printing has taken science and industries by the storm, offering in-house design customization, fast iterative development, and cheap production using professional printers based on filament extrusion (e.g., MakerBot Industries, New York, NY) and liquid resin (e.g., Form 3 by Formlabs, Sommerville, MA). Yet, plastic prints have limitations mostly due to the low strength of the materials. Recently metal printing has become affordable offering increased strength, with options for printing in aluminum, stainless steel and even titanium with similar printing resolution to plastics. Here, we have taken advantage of this opportunity and constructed a 3D printed microdrive from stainless steel (stainless steel 316L, 20 μm resolution), which offers superior strength (~10 times higher than plastic) and form factor compared to plastic prints. The metal printing allows for the drives to be reused with minimal wear, driving the effective cost down.

The microdrive is composed of three metal parts: an arm, a body, and a base (Figure 1A and B) and has a footprint of 15.5 mm^2^. The detachable base allows for easy recovery of probes. The arm/shuttle is mounted on a screw to the drive body, allowing it to move linearly along the vertical axis simply by turning the screw (270 μm/ turn). The constructed microdrive has a total travel distance of 6 mm, allowing one to record from distant brain regions across days. Due to its small form factor, multiple probes can be implanted in the same animal (Figure 1C). It comes with a stereotaxic implantation tool for user-friendly and reliable implantations and probe recovery, consisting of a stereotactic manipulator attachment and a microdrive holder (Figure 1D and E).

**Figure 1.**
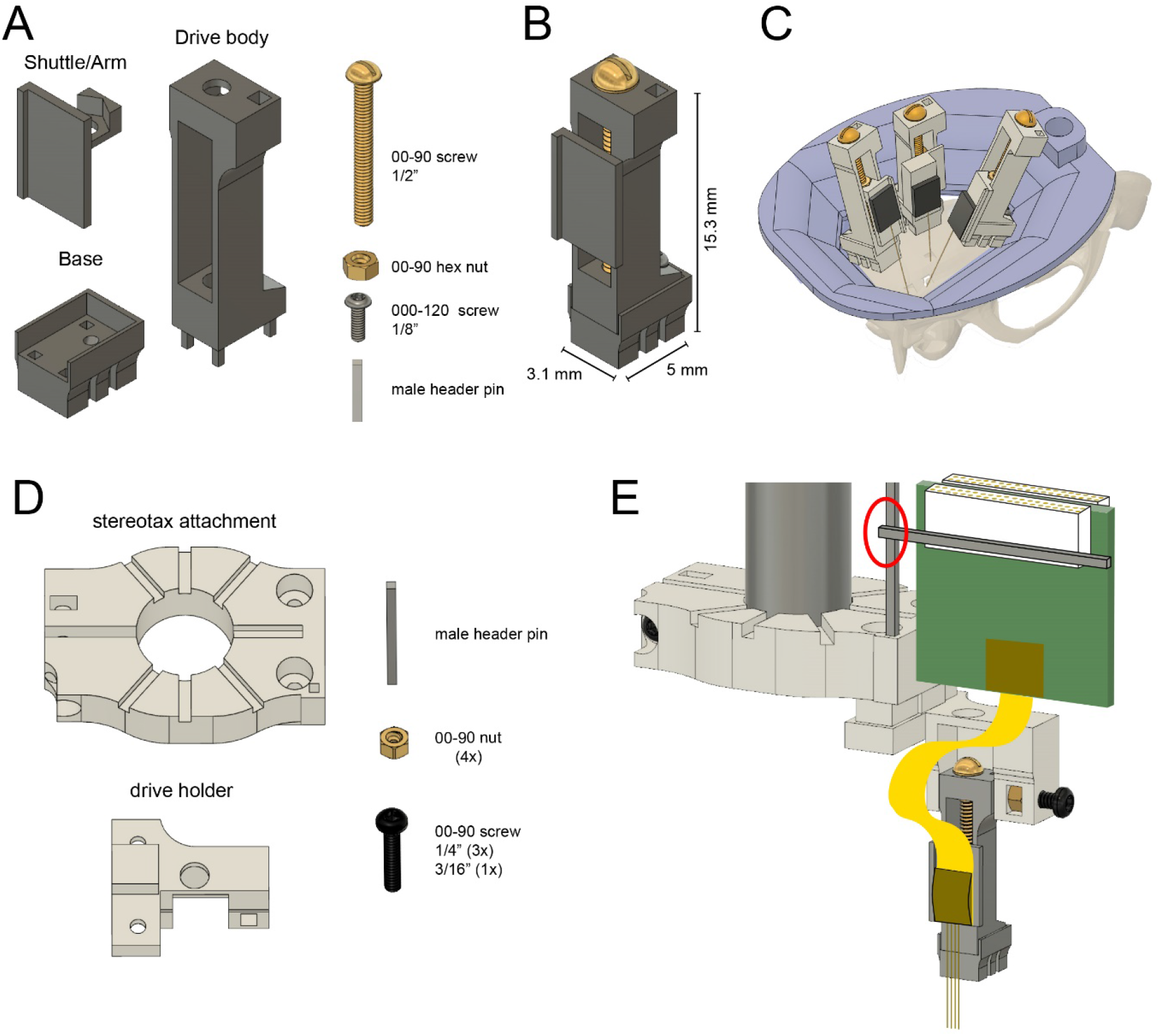
Reusable metal microdrive. **(A)** The metal microdrive consists of three main parts: a drive body, a movable arm/shuttle, and a removable base. All components are 3D printed in stainless steel. Additional necessary components are a 00-90, 1/2 “brass screw, a 00-90 brass hex nut, a 000-120, 1/8” stainless steel screw fixing the drive to the base, and a male header pin. **(B)** The assembled drive with dimensions. **(C)** Schematic showing three microdrives, with silicon probes attached, implanted in a rat to target hippocampus, medial and lateral entorhinal cortices. 3D printed resin head cap is shown in purple. **(D)** 3D printed stereotaxic attachment and drive holder together with assembly pieces: male header pin, four 00-90 brass hex nuts, three 00-90, 1/4” and a 3/16” stainless steel screw. **(E)** Stereotaxic attachment with the metal drive assembled, and a probe attached, ready for implantation (red circle highlights the temporary soldering joint for the Omnetics connector).

The fully assembled microdrive weighs 0.87 g (base: 0.23 g, shuttle/arm with nut: 0.16 g, drive body with screw and metal bar: 0.49 g). This weight and dimensions are similar to other commercially available or custom-made electrode microdrives (Table 1). The design files for our microdrive can be submitted to commercial 3D printing companies (e.g., Proto Labs, Maple Plain, MN) allowing for high-quality printing and fast production. The printing costs of the 3 components are about $140, a highly competitive price compared to commercial microdrives.

### Mouse cap

To make silicon probes truly reusable, both the microdrive and the head cap have to be reusable. The mouse cap is composed of three parts: a base, a left-side wall, and a right-side wall (Figure 2A). The cap-base is attached to the skull of the animal during anesthesia using a ring of Metabond cement, serving as a base for the rest of the cap. There is no need for skull support screws, making the head cap minimally invasive. The cap has a large internal window shaped as an elongated octagon, following the outer ridge of the skull, giving wide access for various surgical needs (Figure 2B). The sidewalls provide structural support, electrical shielding (by acting as a Faraday cage), and physical protection of the silicon probes, hardware, and electronics. The internal volume allows for great flexibility and can fit two Omnetics preamplifier-connectors, as well as optic fibers. The sidewalls attach to the base using a rail and with three support screws.

**Figure 2.**
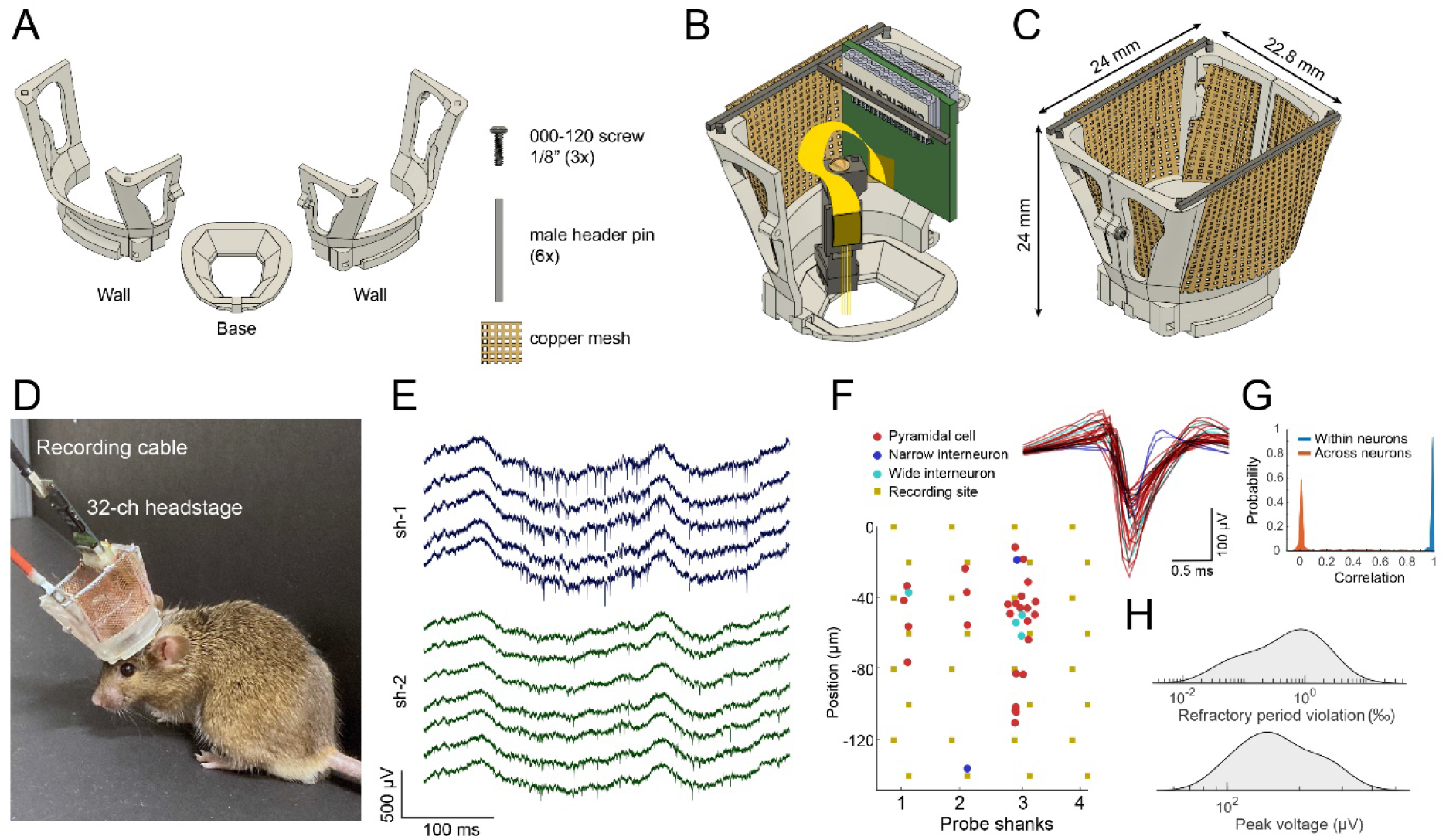
Mouse cap. **(A)** The mouse cap consists of three main 3D printed parts: a base, and two side walls. The pieces are assembled with three 000-120, 1/8” steel screws, six male header pins, and copper mesh. **(B)** The base with the left side wall attached. Copper mesh was attached in three pieces to the wall, and a male header pin was soldered across the top of the wall. **(C)** The fully assembled mouse cap. **(D)** The implanted headgear with preamplifier and recording cable attached. **(E)** Wide-band extracellular traces recorded from the prelimbic cortex of the implanted mouse shown in (**D**) using a multi-shank silicon probe during food pellet chasing exploration (sh-1 and sh-2 denote shank-1 and shank-2 of the silicon probe). **(F)** Well isolated single units can be recorded using the mouse cap system and microdrive (n = 31 putative single units; same session as in E). The location of the maximum waveform amplitude of each neuron is shown on the left side (0 μm corresponds to the location of the topmost channel of the shank. Four shanks are shown). The waveforms are shown on the right (putative pyramidal cells, putative narrow waveform interneurons and putative wide waveform interneurons are shown in red, blue, and cyan, respectively). **(G)** Waveform correlations for each single unit comparing waveforms between the first and the second half of the recording within the same neuron (blue) and between neuron pairs (red). **(H)** The refractory period violations and the peak voltage across the recorded neurons (n = 31).

The entire cap weighs 2.2 g (base: 0.19 g, walls with male header pins and copper mesh: 0.98 g each, and 000-120 screws: 0.05 g; Figure 2C). A chronically implanted mouse can carry this cap with one (or more) implanted silicon probe and with a custom connector for electrical stimulation (Figure 2D). High-quality electrophysiological signals can be collected from freely moving mice for weeks and months (Figure 2E and F). The system can be customized further, using our CAD files (see Methods section). We recommend printing the cap system on the Formlabs Form 2/3 resin printer or a comparable 3D printer (requires 25-50μm resolution).

### Rat cap

The typical Long-Evans rat is approximately ten times heavier than the mouse (~400g), and requires a sturdier cap system, capable of withstanding forceful impacts and provide increased protection of the electronics and hardware. The rat cap is composed of four parts: a base, a left-side wall, a right-side wall, and a top cover (Figure 3A). The octagon-shaped base aligns with the outer rim of the rat’s dorsal skull surface and is attached with Metabond cement, with no need for skull support screws, making it minimally invasive (Suppl. Figure 1A). The two side walls are attached to the base with a single rotation-axis located in the front of the base, attached with a long screw (Figure 3B, top part). The walls are held in place on the base, using a rail and two screws in the back. The sidewalls have two sets of male header pins for soldering standard Omnetics probe connectors (see Surgical Instructions). The lid can be locked with a thumb screw and has holes for air ventilation (Figure 3B bottom part). High-quality electrophysiological signals can be collected from freely moving rats for weeks (Figure 3C and D).

**Figure 3.**
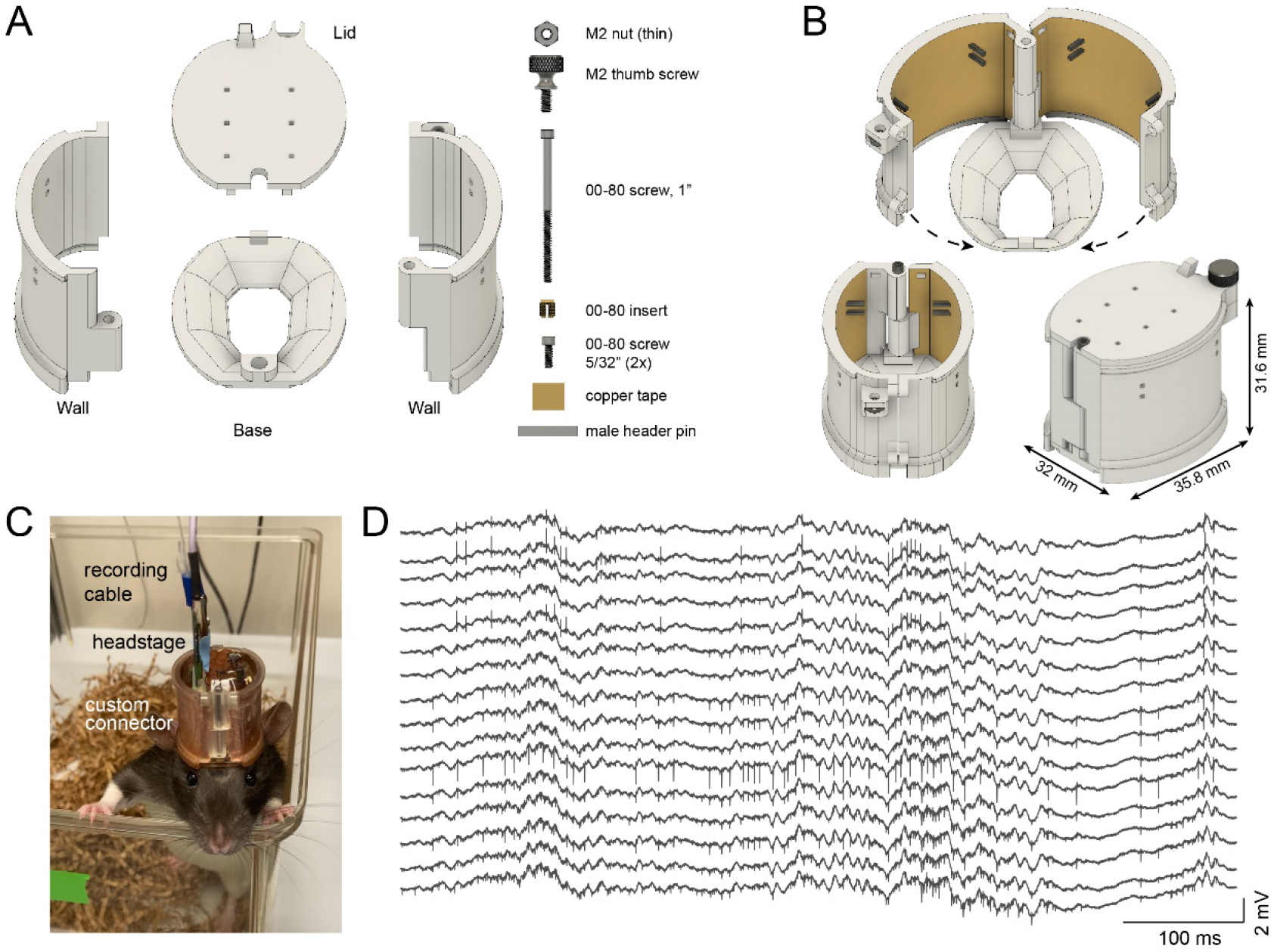
Rat cap. **(A)** The rat cap consists of four main 3D printed plastic parts: a base, two side walls, and a lid. To assemble the components, an M2 nut, M2 thumb screw, a 00-80, 1” screw, a 00-80 insert, and two 00-80, 5/32” screws are also needed. **(B)** The assembled rat cap is shown with sidewalls in an open position (top image), closed configuration without (bottom left) and with the lid in place (bottom right). **(C)** A Long-Evans rat in its home cage with the rat cap, connected to preamplifier and cable. **(D)** Extracellular traces from the same animal.

For complicated experiments, the cap system can be modified to increase the available skull surface (Suppl. Figure 1A and B). This modified base is held by bone screws implanted in the temporal bone and covered with dental cement (Suppl. Figure 1B right part). Increasing the inner volume of the cap system and using metal recoverable microdrives enable multiprobe implantations (Suppl. Figure 1C).

The entire design weighs 11.03 g (base: 1.04 g, right wall with male header pins and copper tape: 3.48 g, left wall with male header pins and copper tape: 3.68 g, top with thumb screw: 2.35 g and 00-80 screws: 0.48 g; Figure 3B bottom, right).

### Surgical advantages using the head cap systems

The modular system decreases the duration of the surgery and allows for faster post-operative recovery for the animal, due to four important modifications. 1. The head cap is prepared before surgery and can be reused easily. 2. The cap does not need for support screws, reducing the invasiveness of the surgery and accelerating the animal’s recovery. 3. The 3D printed cap-base is secured with a single step, by attaching it to the dorsal surface of the skull with Metabond cement. This ensures alignment precision relative to the brain surface, easier probe recovery, and reusability. 4. The electric shielding and structural support is implemented in the reusable head cap, decreasing extra manual steps for the construction of the protective cap from copper mesh, male header pins and grip cement during surgery (Vandecasteele et al., 2012).

These steps offer a time savings from 40 to 90 min (Suppl. video 1A and B), compared to a manually constructed cap during surgery (Vandecasteele et al., 2012).

Further, the modular cap system substantially increases flexibility during an implantation procedure. Because the sides can easily be disassembled and reassembled, a complex surgical produce can be split into multiple sessions when needed. In the first session the skull is prepared, and the base of the cap is attached to the skull. After recovery, the craniotomy and implantation are performed in a second surgery. This result in a speedy recovery of the animal and reduces the likelihood of human error during the procedure. Additionally, subsequent troubleshooting can be made throughout the chronic experiment with minimal disruption to the animal and the implanted devices.

### Probe recovery

To recover the probe at the end of the chronic experiment, the drive holder is aligned with the drive using the stereotactic frame. Once the position is aligned in the x-y plane, the drive holder is moved downwards (Figure 4A, step 1). Next, the top of the drive is secured with the screw located on the side of the drive holder (Figure 4A, step 2). The 000-120 screw is removed from the base (Figure 4B, step 1) and the drive is moved upwards carefully (Figure 4B step 2 and C). It is recommended to monitor the shanks of the probe under a microscope during the entire recovery procedure and, if any unexpected movement of the probe is observed, return to the previous step to make sure that everything is secured properly.

**Figure 4.**
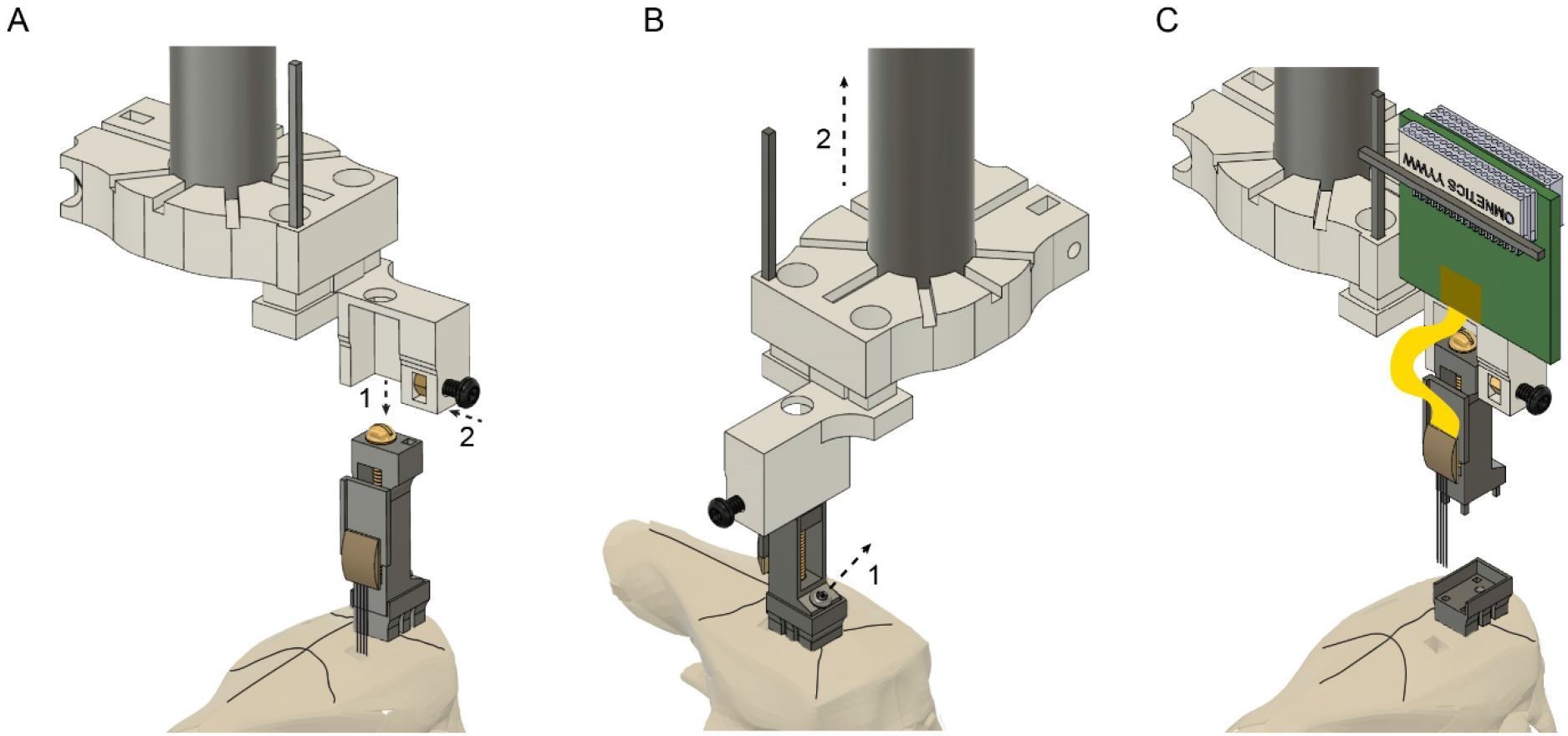
Probe recovery procedure. **(A)** The stereotaxic probe holder is attached to the microdrive (step 1) and is fixed with the black screw (step 2). Precise alignment is critical to avoid tissue damage and prevent breaking the probe shanks when retracting the probe. **(B)** The microdrive is detached from the drive-base by removing the 000-120 steel screw (step 1) and moved upwards (step 2). Camera angle rotated 90°. **(C)** The drive with the attached probe after retracting it from the brain. The drive-base can be reused by cleaning it in chloroform or acetone.

The removed probe is cleaned by initially rinsing it in distilled water, then contact lens solution (containing protease) and distilled water again; each washing step should last for at least 12 hours. If extra tissue or debris is detected between the shanks, it can be carefully removed by a fine needle (26 gauge or smaller) under a microscope.

## DISCUSSION

We have developed a recoverable microdrive printed in stainless steel and a head cap system for chronic electrophysiological recordings in freely behaving rats and mice. The cap system allows for considerably faster and more standardized surgeries to be performed and faster post-surgical recovery of the animals. Importantly, recovery of the probe and head cap becomes an easy and routine procedure, allowing the same silicon probes to be used in multiple animals, offering substantial savings.

Our head caps are minimally invasive and do not require bone screws. Except for the base, the entire head gear is reusable, making experiments performed on multiple animals less variable. For multiple surgeries (e.g., virus injection for optogenetic or pharmacogenetic experiments), implantation of the base during the first surgery provides fixed coordinates for a subsequent surgery. The head cap system is flexible, due to the large internal volume, and allows for multiple probe implants, optical fiber implants and other optional components. In contrast, manually constructed cap systems are time consuming to build, require extensive experience and its construction may vary from animal to animal and across investigators even in the same laboratory. The main disadvantage of hand-built head gears is the limited success for probe recovery. Even after successful recovery of the recording probe, a new protective cap must be built from scratch in future surgeries. In contrast, our modular cap system is prepared before surgery, decreasing the time the animal spends under anesthesia, reducing potential complications during and after surgery. Using this strategy, we were able to explant and implant the same silicon probe in >10 mice (Senzai et al., 2019).

The metal microdrive weighs 0.87 grams with a footprint area of 15.5mm^2^, allowing the implantation of multiple probes in rats, and even in mice. Because the entire headgear can be removed from the base with a screwdriver, recovery of the silicon probes is simple and highly successful. The drives are printed in stainless steel, with a stiffness (Young’s modulus) approximately ten times higher than that of plastic. Steel drives provide higher stability, better recording quality and prevent potential wobbling while turning the screw to adjust the probe’s position in the brain. Commercially available drives are typically built from plastic, they are non-recoverable and more expensive. Hand-made drives introduce variability across drives and experiments. In contrast, 3-D steel printing provides high consistency across drives, reducing interexperimental variability.

To facilitate wide use of the 3D printed designs, we share all necessary details of parts, fabrication process and vendor source for easy replication by other laboratories. We offer several video tutorials, which describe the construction of the microdrive, the cap systems, the probe implantation, and the probe recovery. The CAD system allows different laboratories to customize both the drive and headgear according to their specific goals and needs.

## MATERIALS AND METHODS

### Key resource table

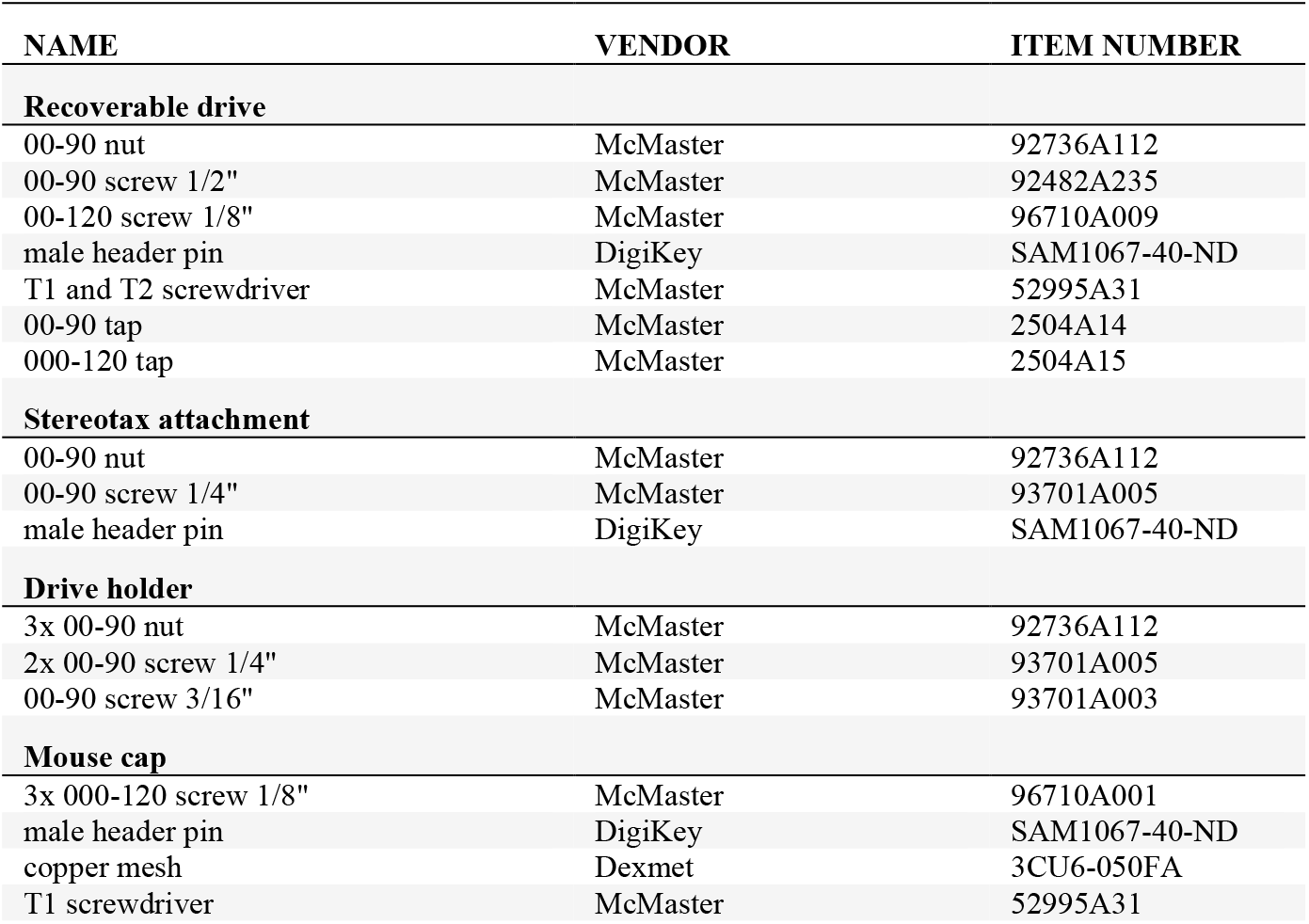
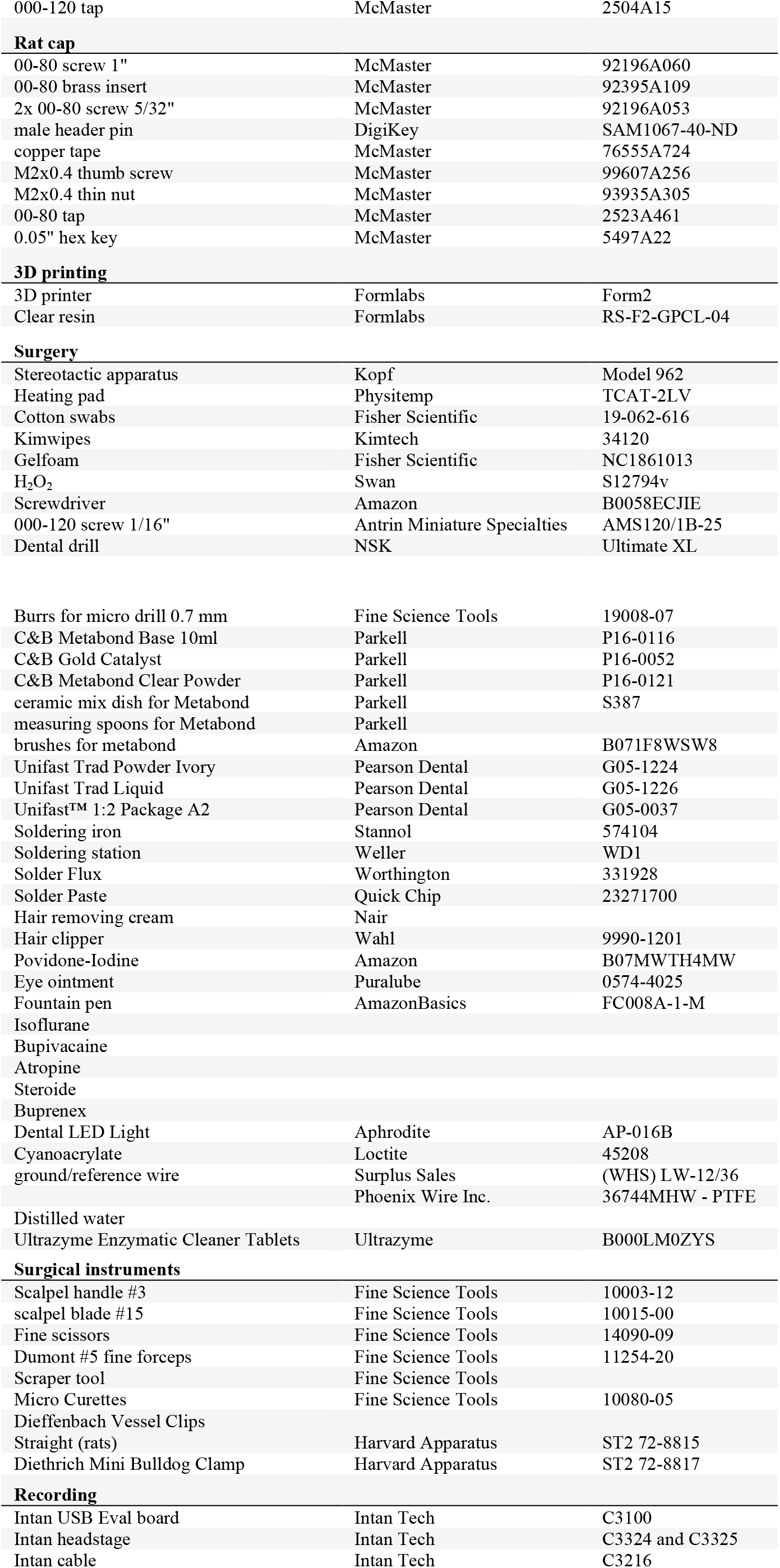

### Microdrive assembly instructions

The base of the microdrive provides an anchoring point for the body of the microdrive via a tapped hole in the back (000-120 tap) and rectangular holes inside the base (0.5 × 0.5 mm). Thin walls around the base prevent cement flowing between the base and the body during surgery (Figure 1A). Glue a nut inside the arm (referred to as ‘arm nut’; 00-90 brass nut) before attaching it to the body. The body has an opening in the top part of the back where a nut can fit inside (‘top nut’; 00-90 brass nut). Insert the ‘top nut’ from the back, then insert the arm from the front and introduce a screw (00-90, 1/2”, brass screw) through the ‘top nut’ and the ‘arm nut’. Tighten the screw completely and release it a quarter-turn (or less). Fix the ‘top nut’ and the screw together using solder so the arm can be moved linearly relative to the body by turning this screw. Attach the body-arm complex to the base using a screw in the back (000-120, 1/8”, stainless steel screw). Finally, insert a male header pin into the body and secure it using dental acrylic cement (Unifast Trad). This can be used as a soldering joint during surgery. Finally, attach the backend of the silicon probe to the arm using cyanoacrylate glue and solder the Omnetics connector (Omnetics Connector Corporation) of the probe to the male header pin of the drive holder. The fully assembled microdrive weighs 0.87 g (base: 0.225 g, arm with nut: 0.159 g, body with screw and metal bar: 0.486 g).

*Assembly_instructions_microdrive_metal_v7.pdf* contains instructions with photographic documentation.

### Implantation/recovery tool assembly instructions

Insert and glue one nut (00-90, brass nut) and a male header pin into the stereotactic attachment using cyanoacrylate glue. Insert a 00-90, 1/4“ stainless steel screw into the nut. Tightening this screw will secure this piece to the electrode holder of the stereotactic arm (Model 1770, Kopf Instruments). The male header pin should be used as a temporary soldering joint for the Omnetics connector of the silicon probe. Insert and glue two nuts (00-90, brass nut) into the bottom of the drive holder and one nut (00-90, brass nut) into the body of the drive holder. Insert a 00-90, 3/16“ stainless steel screw through this latter nut. This screw should be used to secure the top part of the body of the drive to the drive holder. Attach the stereotaxic attachment to the drive holder using 00-90 screws (00-90, 1/4”, T2 screw).

*Assembly_instructions_implantation_tool_metal_v7.pdf* contains instructions with pictures.

### Mouse cap assembly instructions

The base has a rectangular hole for a male header pin (0.8 x 0.8 mm) for fixing the left and right walls temporarily during surgery (Figure 2B). This can help to open the cap using a fine pair of tweezers. The tip of the tweezer is squeezed between the rectangle and the walls. Pushing the tweezer against this rectangle readily opens the walls. The right wall has one tapped hole in the front and one in the lower part of the back (000-120 thread, 1.9 mm length). In addition, it has a hole in the upper part of the back (1 mm in diameter, 1.4 mm length). The left wall has one hole in the front and one in the lower side of the back (1 mm in diameter, 1.4 mm length) and a tapped hole in the upper side of the back (000-120 thread, 1.9 mm length). In addition, there are two rectangular holes in each wall (0.8 by 0.8 mm) in which male header pins are glued with cyanoacrylate glue to serve as soldering points for the Omnetics connector and for the shielding copper mesh. To reduce weight, walls are perforated and covered with light copper mesh by gluing it with dental acrylic (Unifast Trad). The walls are closed using two screws in the back and one screw in the front (000-120, 1/8” stainless steel pan head torx screws).

*Assembly_instructions_mouse_hat_10_39mm_v11.pdf* file contains instructions with pictures.

### Rat cap assembly instructions

The base has a hole for a brass screw-to-expand insert (00-80 thread size, 1/8" installed length) and serves to hold together the left and right walls. It has a rectangular protrusion in the back (3 by 1.5 by 1.67 mm) to help opening of the cap using a fine tweezer. The right and left walls have a front hole (diameter 1.8 mm) in which a screw can be passed (00-80, 1” 18-8 stainless steel socket head screw) for fixing the walls to the metal insert of the base. In addition, there is a rail on each wall at the bottom part that grabs onto the base piece (1.2 mm height and 1 mm deep).

During surgery, the walls are kept open with the screw loosely tightened (Figure 3B top part). After all the connectors are attached to the male header pins, the walls are closed, and the front screw is tightened. The right wall has a hole in the upper side of the back (diameter 1.8 mm, 2 mm length) and a tapped hole in the lower side of the back (00-80 thread, 2 mm length). The left wall has a hole in the lower side of the back (diameter 1.8 mm, 2 mm length) and a tapped hole in the upper side of the back (00-80 thread, 2 mm length). The walls are closed in the back using two screws (18-8 stainless steel socket head screw, diameter 0-80, 5/32” length). The left wall also has an insert in the upper part of the back side for a nut (18-8 stainless steel thin hex nut, M2.5 × 0.45 mm thread). This latter nut serves as a locking mechanism for the top cover. There are four rectangular through-holes in each wall (0.8 × 0.8 mm) in which male header pins are glued with epoxy (Araldite® Standard Epoxy) and serve as soldering points. The locations of the holes can be modified according to user specifications to adapt different connector placements. To protect the implanted electrodes, the rat cap is covered by either self-adherent wrap (3M™ Coban™) or the plastic top cover. The edge is extruded on the outer surface on top of the walls to provide extra surface for better adhesion. The plastic cover is attached to the walls using the front slide-in slot and the back screw (stainless steel flared-collar knurled-head thumb screw, M2 × 0.40 mm thread size, 4 mm long). To protect the neuronal signal from environmental electromagnetic interference noise, conductive copper coil electrical tape is glued to the walls by cyanoacrylate glue (copper tape: 1" wide, McMaster product number: 76555A724) and connected to the ground.

*Assembly_instructions_rat_cap_v8.pdf* file contains instructions with photographs.

### 3D designing and printing parts

All parts were designed in Autodesk Fusion 360 (https://www.autodesk.com/products/fusion-360). We tested and printed cap designs on a Form 2 printer from Formlabs with 50 μm resolution using their standard resins. The metal microdrive prints were produced by Proto Labs (https://www.protolabs.com/services/3d-printing/direct-metal-laser-sintering). All design files are available on our GitHub resource (https://github.com/buzsakilab/3d_print_designs).

### Subjects

Rats (adult male Long-Evans, 250-450 g, 3-6 months old, n = 7) and mice (adult male C57BL/6JxFVB mice, 32-40 g, n = 5) were kept in a vivarium on a 12-hour light/dark cycle and were housed 2 per cage before surgery and individually after it. All experiments were approved by the Institutional Animal Care and Use Committee at New York University Medical Center. Animals were handled daily and accommodated to the experimenter before the surgery and behavioral recording.

### Surgery instructions

The following instructions cover surgeries in both rats and mice, with differences highlighted. Prior to surgery, prepare the 3D printed cap, the microdrive(s), the implantation tool and attach a silicon probe to the microdrive (as described above).

We recommend measuring the impedance of the silicon probe before implantation. Lower the probe into 0.9% saline and ground the saline to the recording preamplifier ground. Connect the probe to an Intan preamplifier headstage and to the main Intan board to perform the impedance test.

#### 1. Prepare the stereotaxic apparatus and tools

1. Place the heating pad under the position of the ear bars.
2. Sterilize surgical instruments.
3. Weigh the animal subject.
4. Place all components in alcohol for disinfection.
5. Mice: prepare bupivacaine in an insulin syringe (0.4 - 0.8 ml/kg of a 0.25% solution).

#### 2. Surgery

##### Anesthesia and pre-incision preparations

1. The animal is anesthetized for 3 min (until after it loses its righting reflex) in an anesthesia-bucket with 2.5:1.5 (Anesthetic % to Airflow ratio).
2. Apply a local anesthetic to the tips of the ear bars before insertion (LMX-4 Lidocaine 4% topical cream). Fix the head with ear bars and attach the closed ventilation nosepiece. Once the animal is in the stereotactic apparatus, the level of anesthesia is lowered (1.2 – 2%).
3. Remove the hair above the planned surgery site using either Nair-hair remover or a hair trimmer.
4. Clean the hairless skin with the antiseptic solution and repeat the process two more times (Povidone-Iodine – 10% topical solution). Apply the antiseptic solution with Kimtech wipes using anterior to posterior swipes. The last swipe must be done in one stroke to minimize infections. Between each swipe with the antiseptic solution, the skin is cleaned by 70% alcohol applied with the same technique.

##### Incision and skull cleaning

5. Inject bupivacaine (0.4 - 0.8 ml/kg of a 0.25% solution) subcutaneously along the scalp midline for local anesthesia. Make one injection site and distribute the anesthetics along the midline.
6. Make a median incision from the level of the eyes to the back of the skull (neck).
7. Separate the skin from the skull, pull the skin sidewise and attach four bulldog clips to create a rectangular shape opening. The bulldog clips should be attached to the subcutaneous soft tissue, not the skin.
8. Scrape the skull with a scalpel and remove the periosteum from the top flat surface of the skull. This is necessary to achieve a strong bond with the 3D printed base.
9. Clean the skull surface with saline and vacuum suction.
10. Clean the skull with hydrogen peroxide and rinse it with saline. The hydrogen peroxide is applied with cotton swabs (about 5 seconds) and rinsed quickly thereafter thoroughly with saline. Avoid touching the skin and muscle with the solution.
11. Cauterize any bleedings along the skull and exposed skin.

##### Attaching the base to the skull

12. Prepare the Metabond on ice. Mix four drops of base with 1 drop of catalyzer.
13. Paint, using a brush, the whole surface of the cleaned and dried skull and let it dry.
14. Mix a new solution of Metabond with powder: four drops of base, one drop of catalyzer and 2 scoops of powder and apply a second layer of Metabond paint to the skull surface. Paint also along the edge of the skull surface.
15. Paint the bottom surface of the 3D printed base with Metabond and align it above the skull and attach it to the skull before it solidifies.
16. Paint with Metabond along the inner contact line between the hat base and the skull and create a sealed area inside the hat.
17. Gently hold the hat base in place (for about for 60 seconds) until it stays attached to the skull using your fingers. Let the Metabond cure before proceeding to the next steps.

##### Craniotomy marking and screw placement

18. Align Bregma and Lambda in the same horizontal plane. Determine the position of Bregma using stereotactic coordinates with a fine needle attached to the stereotactic arm.
19. Calculate the relative positions of the probe incision points.
20. Mark the positions of the planned craniotomies with a scalpel (gently make two orthogonal lines crossing at the planned incision points with the scalpel) and a pen (fill the scalpel-drawn lines with the pen).
21. Mark the position of the reference and ground screws with the scalpel/pen.
22. Remove the stereotactic arm.
23. Drill holes for ground and reference screws in the skull above the cerebellum with a high-speed drill. If bleeding occurs, rinse it with saline and vacuum suction until the bleeding stops.
24. Insert the ground and reference screws in. Begin with a slight counterclockwise turn. For mice, allow a margin of about 0.5 mm. In rats, drive the screws tight. Alternatively, 125 μm stainless steel wires can be used for reference and ground, instead of screws.

##### Craniotomy

25. Perform the craniotomy with a high-speed drill (drill bit size depends on the goal). Rinse it with saline and vacuum suction to ensure visibility while drilling.
26. Clean around the craniotomy with the drill or a scraping/sharp scooping tool.
27. Remove the dura with a hook-shaped needle at the planned incision site for probe insertion: bend the tip of the 30G needle to form a small hook (gently tap the tip of the needle into a hard surface to form the hook). Lift the dura with the hook and cut with a pointed scalpel (size 11). Avoid damaging blood vessels.
28. Apply saline and Gelfoam to the craniotomy to maintain a wet brain surface.

##### Probe implantation

29. Place the silicon probe in the implantation tool on the stereotactic arm and position it according to the specified surface coordinates.
30. Lower the silicon probe to the brain surface at the marked coordinates.
31. Insert the probe to the desired target depth in the brain.
32. Fix the base of the microdrive to the skull and hat-base with regular grip cement.
33. Apply silicone to the craniotomy, let the silicone run along the shanks and seal the craniotomy completely. This protects the brain and limits bleedings and blood coagulation. Alternatively, apply a mixture of paraffin oil/wax to the craniotomy with a needle and heat it using the tip of a soldering iron.
34. Solder the reference and ground wires to the corresponding sites on the Omnetics connector.
35. Attach the cap sidewalls to the base.
36. Cover the top with the lid or Coban tape.
37. Turn off the anesthesia and release the animal from the stereotactic setup.

##### Post-operative care

38. Weigh the animal after surgery to determine the weight of the added headgear.
39. Place the animal back in a home cage. The cage should be placed on a heating pad during the first night.
40. Inject Buprenex subcutaneously after 20 min (0.05 - 0.1 mg/kg).

##### General notes

- Apply mineral oil to the eyes of the animal at regular intervals.
- To keep the animal properly hydrated during the postoperative days, provide an aqua-gel and a small container with water. Provide regular rodent pills.

### Additional implantation information

Rats and mice were implanted with silicon probes to record local field potential and spikes from the CA1 pyramidal layer in rats and from the prelimbic cortex from mice. Silicon probes (NeuroNexus, Ann-Arbor, MI and Cambridge Neurotech, Cambridge, UK) were implanted in the dorsal hippocampus (rats: antero-posterior (AP) −3.5 mm from Bregma and 2.5 mm from the midline along the medial-lateral axis (ML); mice: antero-posterior (AP) +1.75 mm from Bregma and 0.75 mm from the midline, 10 degree relative to the sagittal axis). The probes were mounted on a plastic recoverable microdrive to allow precise vertical movement after implantation (github.com/YoonGroupUmich/Microdrive) and implanted by attaching the base of the micro-drives to the skull with dental cement.

After the post-surgical recovery, we moved the probes gradually in 50 μm to 150 μm steps until the tips reached the pyramidal layer of the CA1 region of the hippocampus. The pyramidal layer of the CA1 region was identified by physiological markers: increased unit activity and the presence of ripple oscillations (Mizuseki et al., 2011). In mice, the probe was implanted 500 μm below the surface of the brain and recordings were performed each day. The probe was moved 70 μm after each recording day. Data was collected daily.

### Electrophysiology data

Electrophysiological recordings were amplified using an Intan recording system: RHD2000 interface board with Intan 32 and 64 channel preamplifiers sampled at 20 kHz (Intan Technologies). All data is available from https://buzsakilab.com/wp/database/ (Peter Christian Petersen et al., 2020).

### Spike Sorting and data processing

Spike sorting was performed semi-automatically with KiloSort (Pachitariu et al., 2016) github.com/cortex-lab/KiloSort, using our pipeline KilosortWrapper (a wrapper for KiloSort, github.com/petersenpeter/KilosortWrapper (Peter C. Petersen et al., 2020), followed by manual curation using the software Phy (github.com/kwikteam/phy) and our own designed plugins for phy (github.com/petersenpeter/phy-plugins). Finally, we processed the manually curated spike sorted data with CellExplorer (Petersen and Buzsáki, 2020).

## ACKNOWLEDGEMENTS

We thank Manuel Valero, Antonio Fernandez Ruiz, and Kathryn McClain for useful comments on the manuscript. Supported by U19 NS107616, U19 NS104590, R01 MH122391, and The Lundbeck Foundation.

## Author contributions

MV, PCP, and BV designed and fabricated the drive and head cap, MV performed surgeries, MV and PCP processed data, MV, PCP and GB wrote the manuscript.

## SUPPLEMENTARY MATERIELS

**Supplementary Figure 1.**
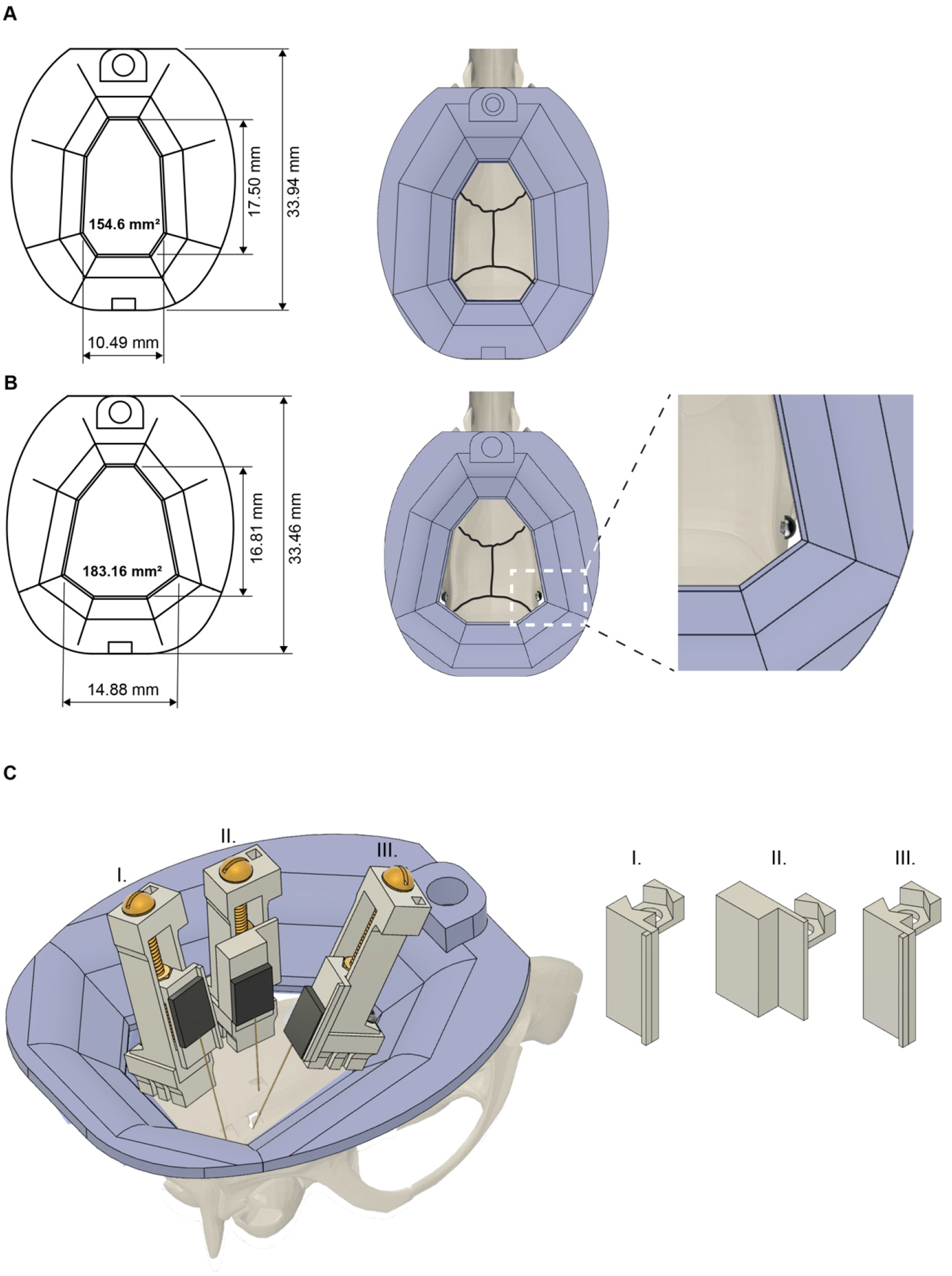
Headcap and microdrive customization. **(A)** Rat cap base with dimensions (left) and on top of a rat skull (right). Black lines indicate the sutures of the skull. 3D printed resin base is shown in purple. **(B)** The base/cap system can be modified to increase the available skull surface (note, that the skull surface was increased to 183.16 mm^2^ from 154.6 mm^2^). This modified base is held in place by bone screws implanted in the temporal bone and covered with dental cement (right part). **(C)** Increasing the inner volume of the cap system and customizing the metal recoverable microdrives enable multiprobe implantations. Schematic showing three microdrives, with silicon probes attached, implanted in a rat to target hippocampus (drive - II), medial and lateral entorhinal cortices (drive - III and I, respectively). Note, the different arm/shuttle designs (right part).

**Supplementary Video 1A. Assembly of the 3D-printed head cap.** 3D-printed head cap is assembled on top of a plastic rat skull. The rat skull with a base is secured in a stereotactic frame (Kopf Instruments). Note the head cap can be assembled completely and closed in less than 2 minutes.

**Supplementary Video 1B. Assembly of the copper mesh head cap.** A protective cap is built on top of a plastic rat skull using copper mesh and dental cement. The rat skull with copper mesh is secured in a stereotactic frame (Kopf Instruments). First, the copper mesh is cut and shaped, then it is covered by dental cement for further mechanical support. Note the head cap can be assembled completely in 30 minutes.

## Notes

### Competing Interest Statement

The authors have declared no competing interest.

