## Supplementary material for "Metal microdrive and head cap system for silicon probe recovery in freely moving rodent": Supplementary_material.docx

#### Recoverable metal microdrive

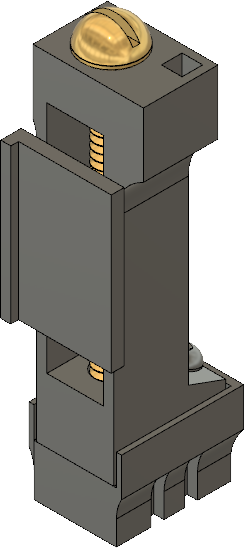
Design name: metal_v7

Travel distance: 5.7 mm Shell base: 3.1 x 5 mm (WxL)

More information is needed:

- contact me (Misi Voroslakos) directly at

arm/shuttle	drive body

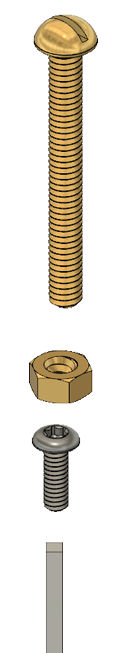
00-90 screw 1/2”

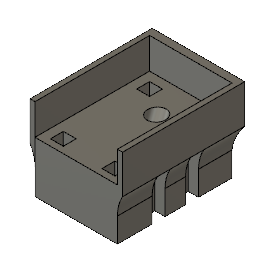

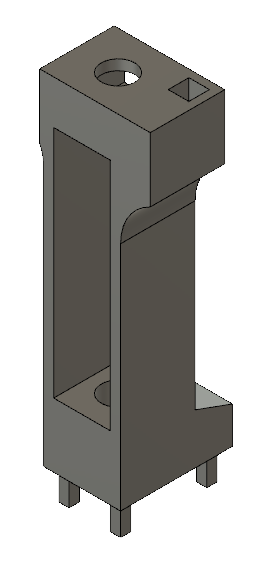

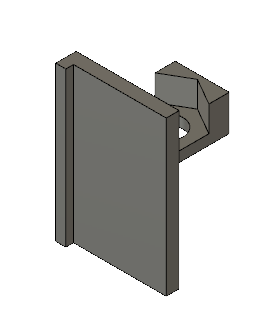

base

brass hex nut

000-120 screw 1/8”

male header pin

#### 1. Tap base (000-120 tap).

COMPLETED STEP

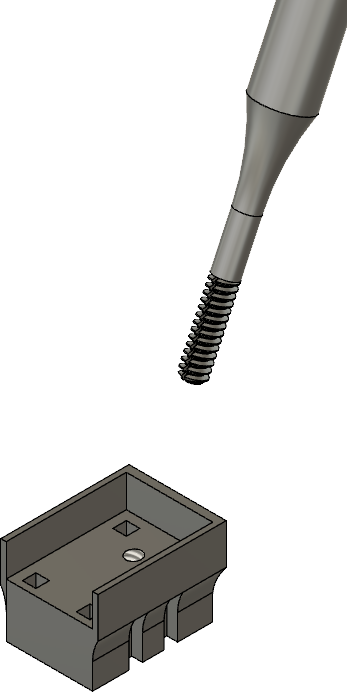

000-120 tap

threaded hole

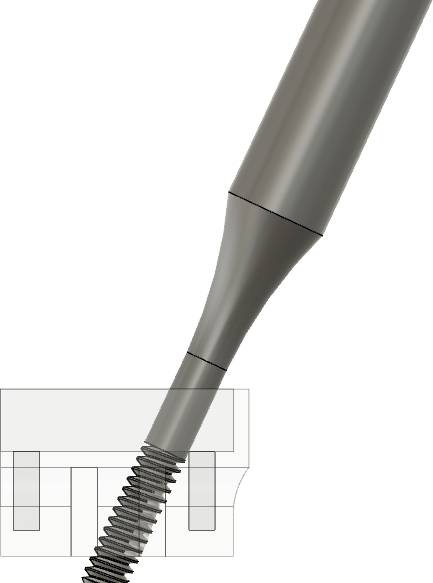

000-120 tap

base

base

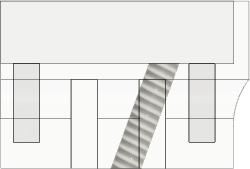

base

#### 2. Insert and glue nut into arm.

00-90 nut	COMPLETED STEP

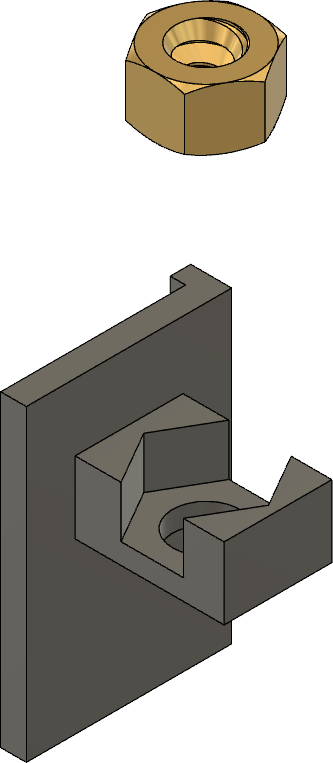

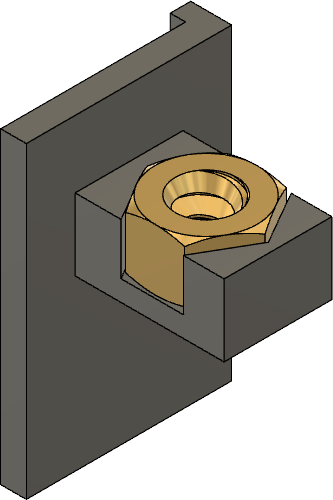
Arm	Arm

#### 3. Solder 00-90 hex nut to 00-90 screw.

**3a - Insert nut into drive.	3b - Insert arm into drive.**

COMPLETED STEP	COMPLETED STEP

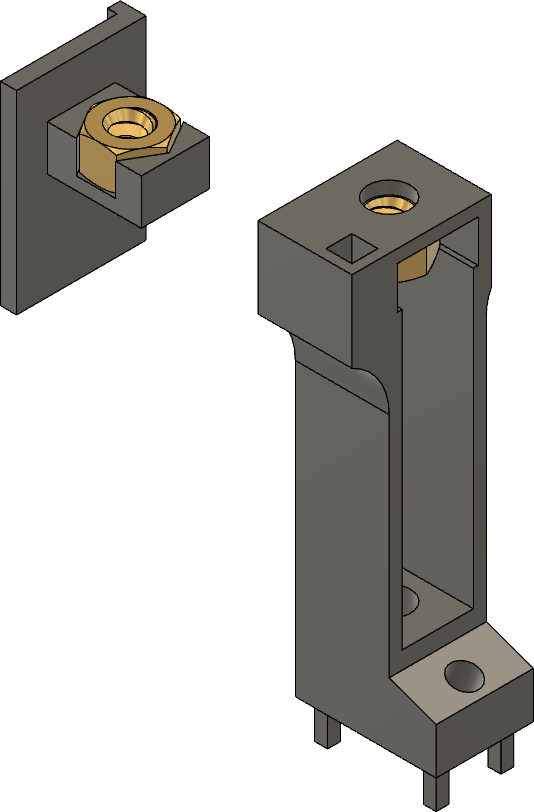

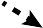

00-90 nut

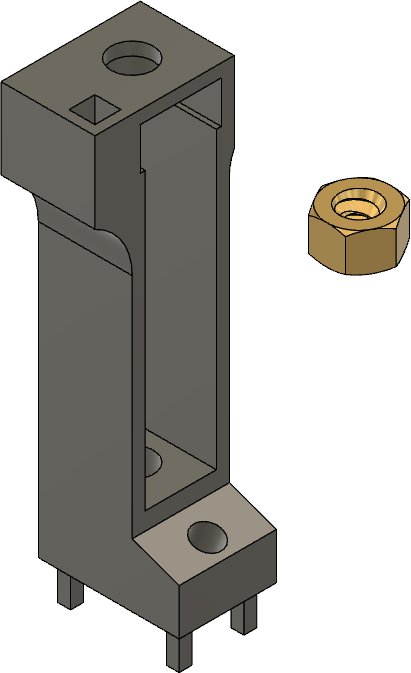

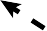

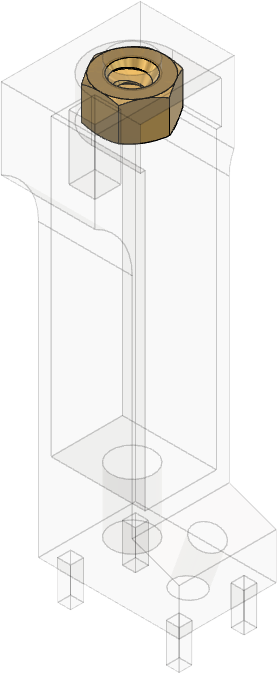
arm

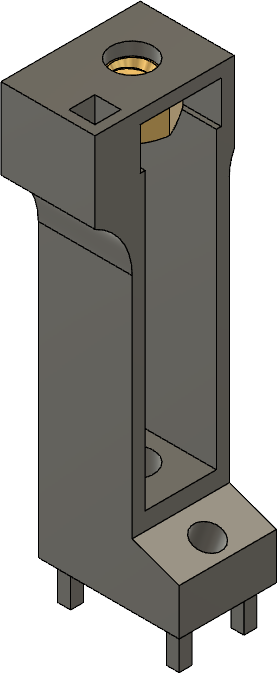

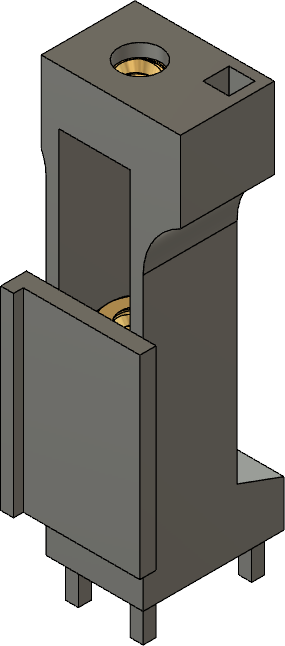

Ke he

arm

drive

00-90 nut

ep gap

re!

drive

drive	drive	drive

#### 3c - Insert 00-90 screw into drive and tighten nut.

00-90 screw

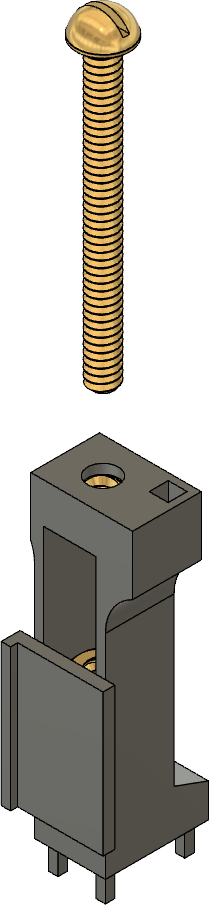

arm

drive

COMPLETED STEP

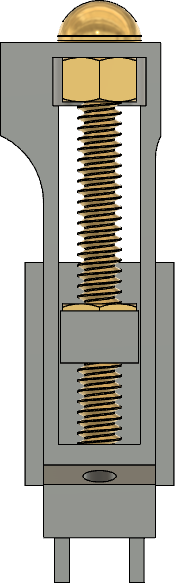
(back view)

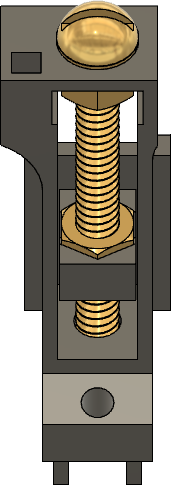

arm

arm

drive	drive

Keep a tiny gap here!

Too tight -> screw won’t move Too loose -> arm will waggle

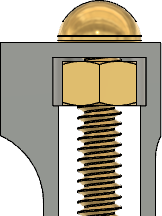

#### 3d - Solder 00-90 nut to 00-90 screw.

00-90 screw

COMPLETED STEP

drive

00-90 nut

drive

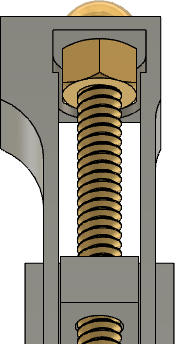

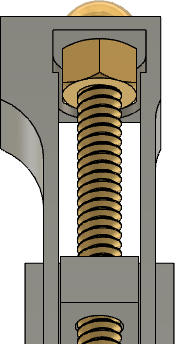

solder here	solder

arm	arm

#### 4. Attach base to drive.

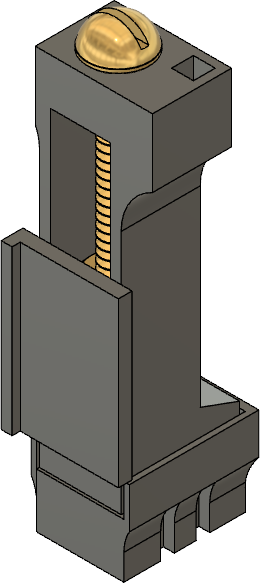
COMPLETED STEP	COMPLETED STEP

drive

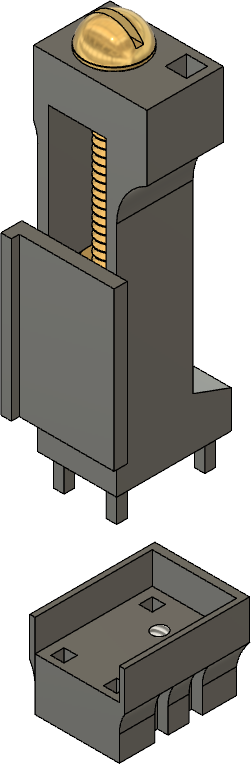

drive

drive

drive

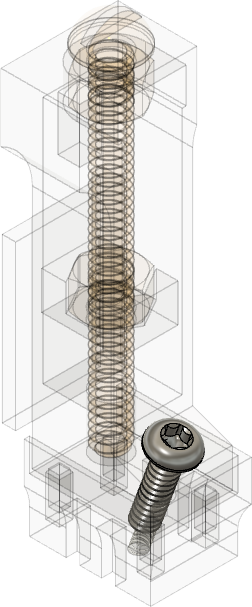

base

arm

arm

arm	arm

000-120

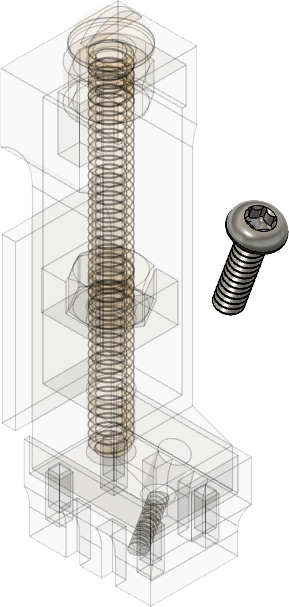

screw (T1)

base

base

base

#### 5. Insert male header pin into drive (if used during surgery).

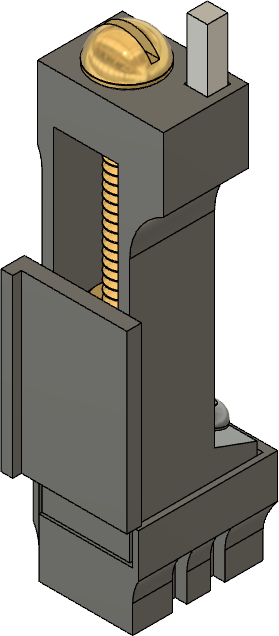
male header pin	COMPLETED STEP

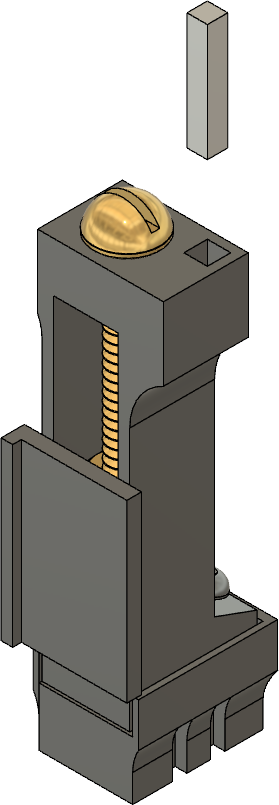

drive

drive

male header pin

arm

arm

base	base

#### Implantation tool for recoverable metal microdrive

Design name: stereotax_attachment_metal_v7

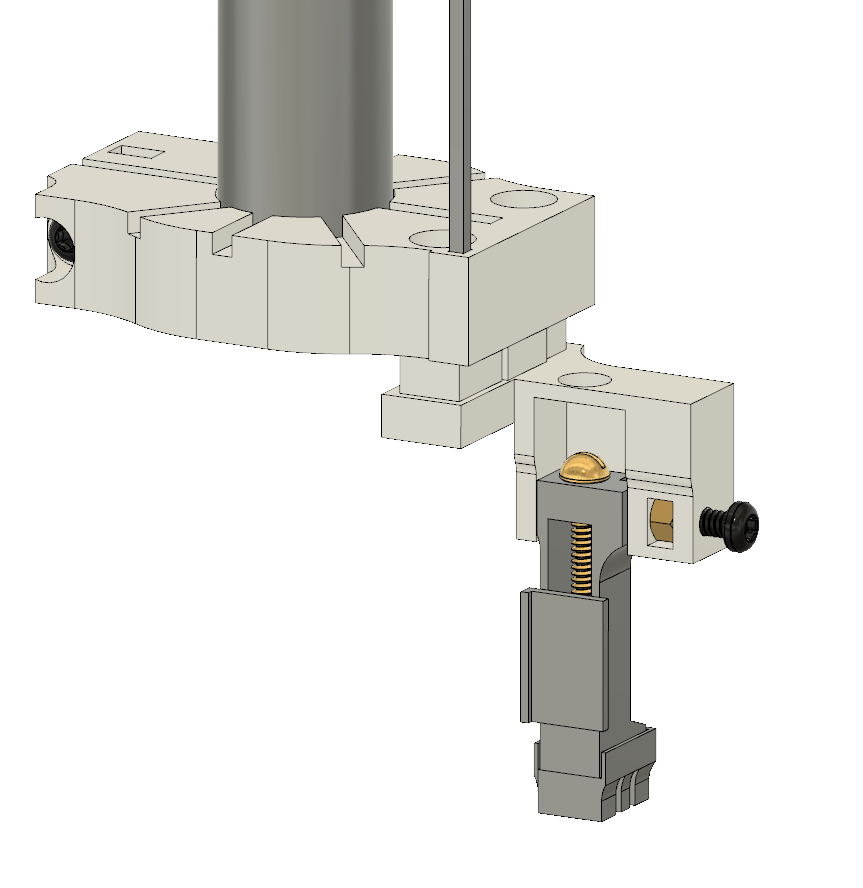

drive_holder_metal_v7

More information is needed:

- contact me (Misi Voroslakos) directly at

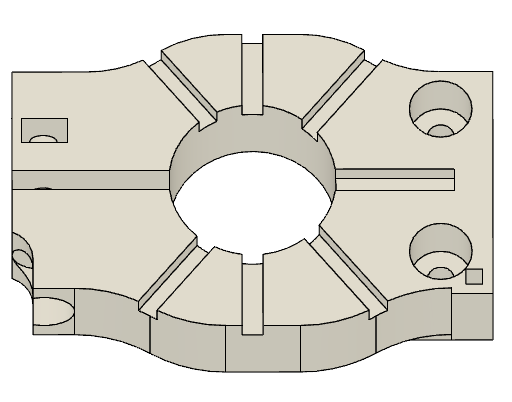
stereotax attachment

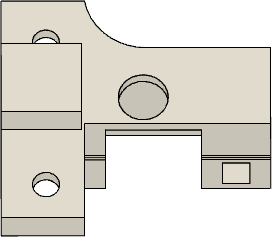
drive holder

### male header pin

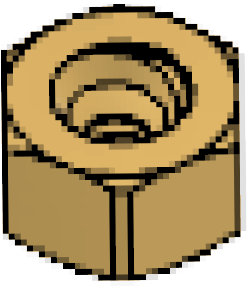

00-90 nut (4x)

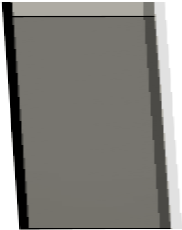

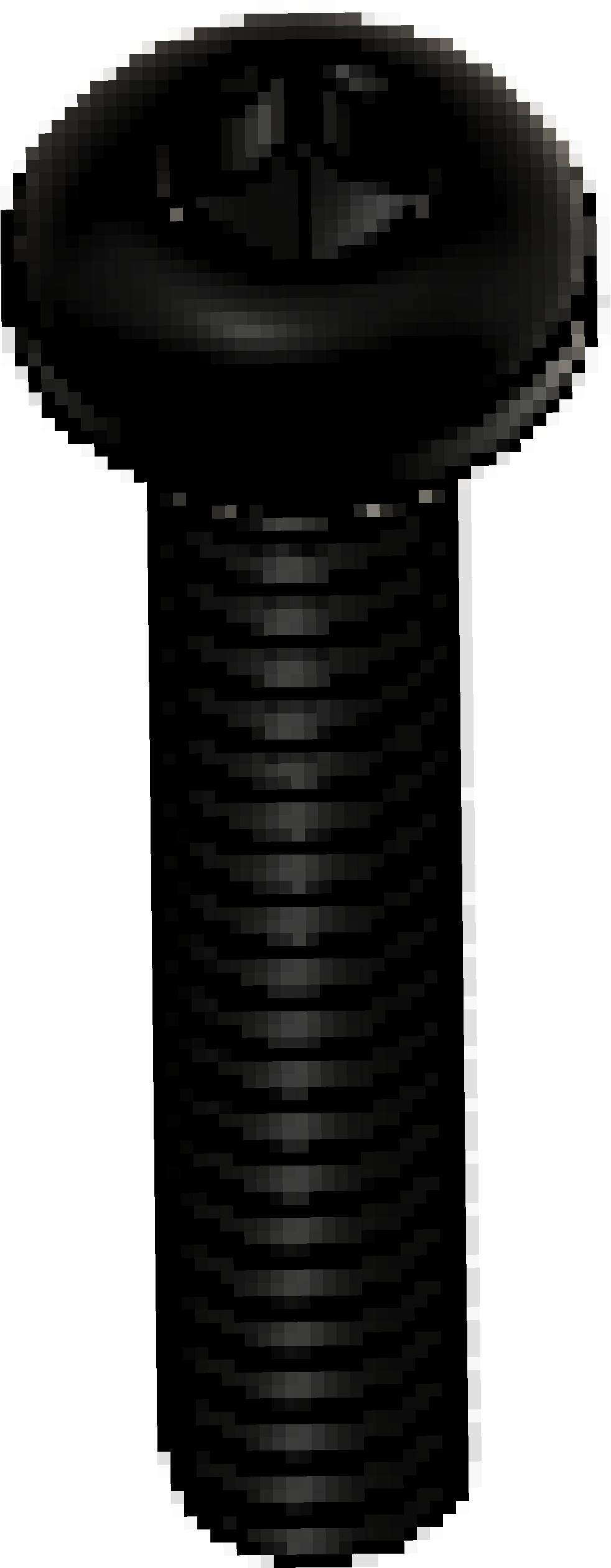
00-90 screw 1/4” (3x)

# 3/16” (1x)

#### 1. Insert and glue 00-90 nut into stereotax attachment.

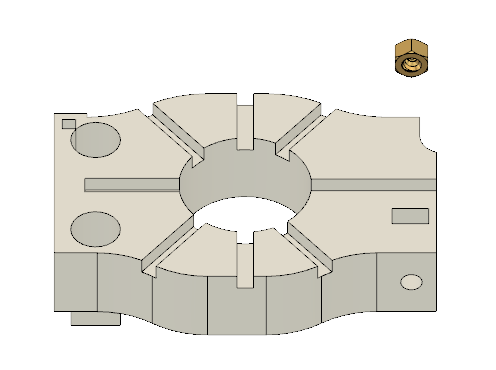

00-90 nut

stereotax attachment

#### 2. Insert and glue male header pin into stereotax attachment.

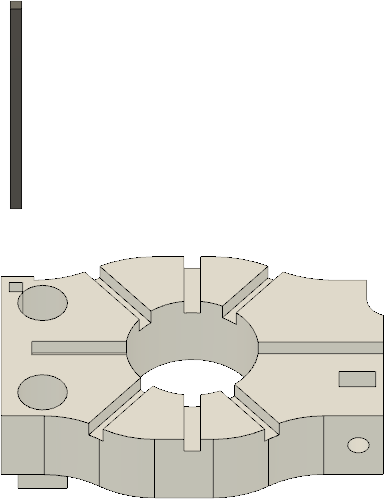

male header pin

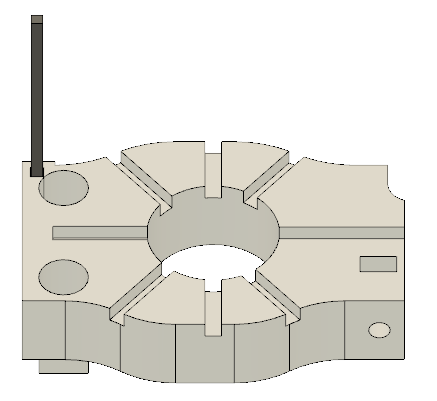

COMPLETED STEP

stereotax attachment	stereotax attachment

#### 3. Insert 00-90 1/4” T2-screw.

COMPLETED STEP

T2

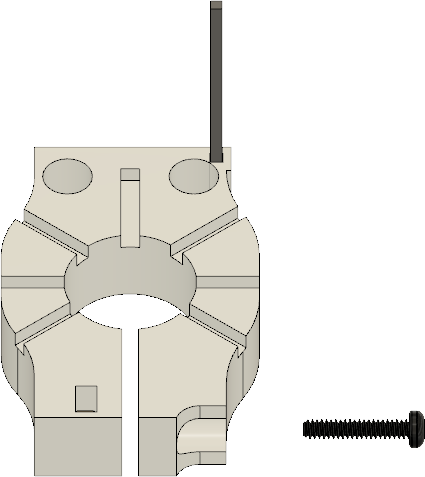

00-90; 1/4”;

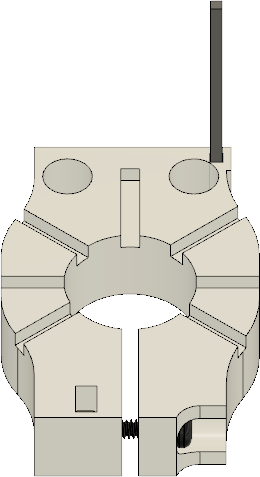

stereotax attachment

pg. 2

#### 4. Insert and glue 00-90 nuts (3x) into drive holder.

00-90 nut

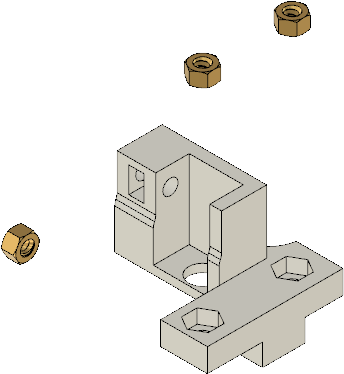

COMPLETED STEP

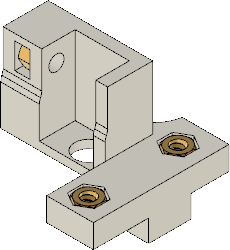

drive holder	drive holder

#### 5. Insert 00-90, 1/4” T2-screws (2x). 6. Insert 00-90, 1/4” T2-screw.

1/4” screw

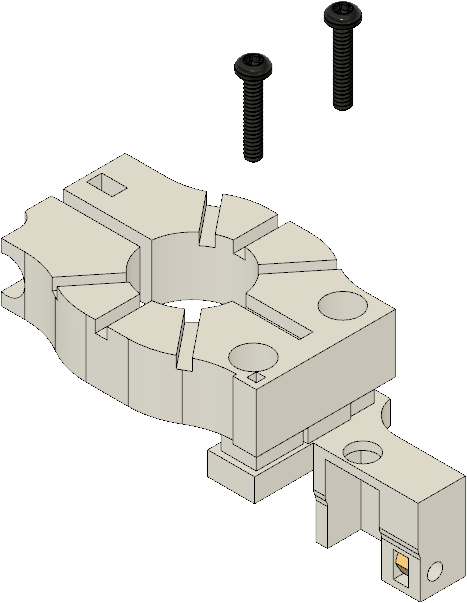

00-90,

stereotax attachment

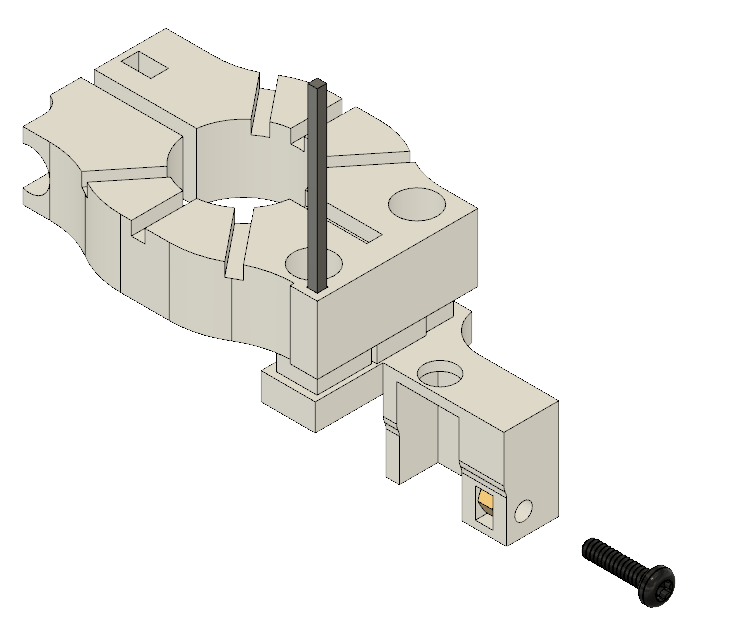

00-90, 1/4” screw

drive holder

pg. 3

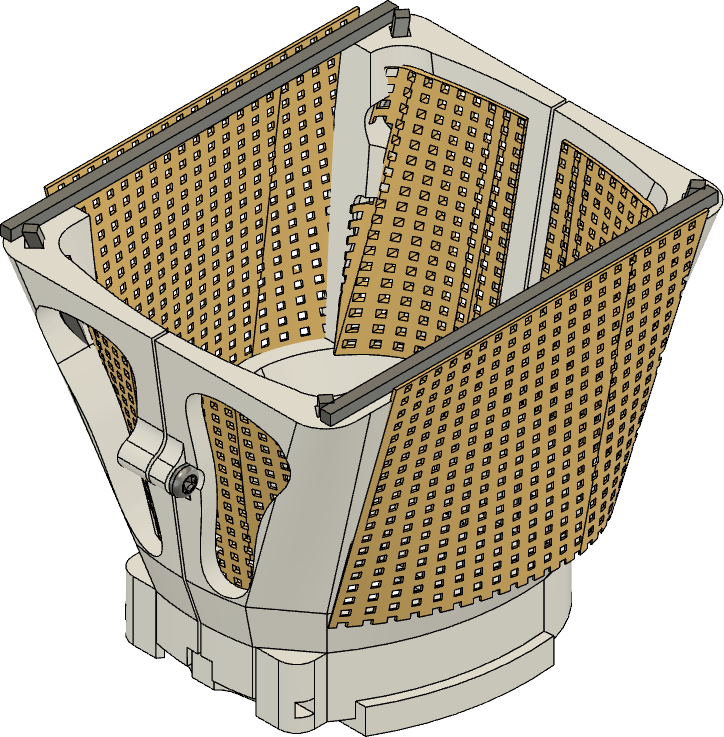
**3D-printed mouse cap**

Design name: mouse_hat_10.39mm_v11 More information is needed:

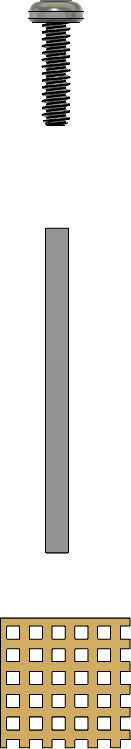
- contact me (Misi Voroslakos) directly at

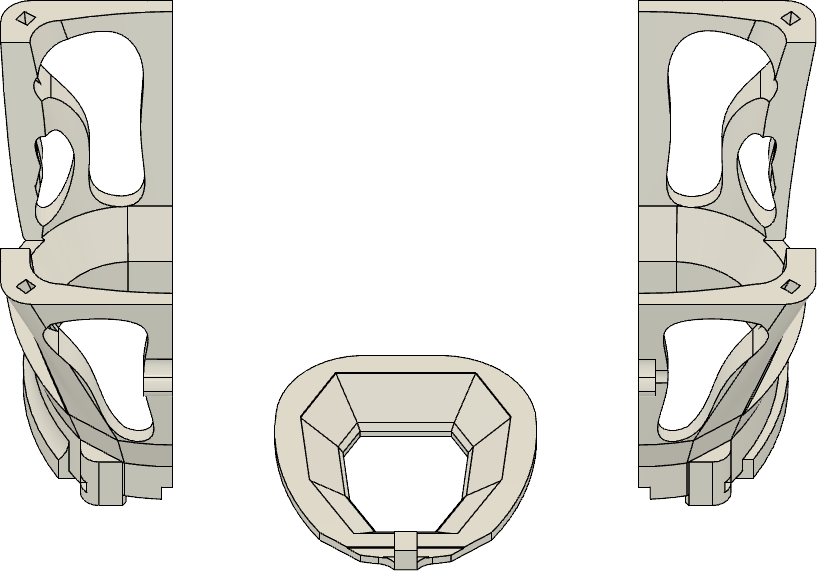

Wall

Wall

Base

00-120 screw 1/8” (3x)

header pin (6x)

copper mesh

#### 1. Tap the walls (00-120 tap).

**1a - Tap the front hole**	COMPLETED STEP

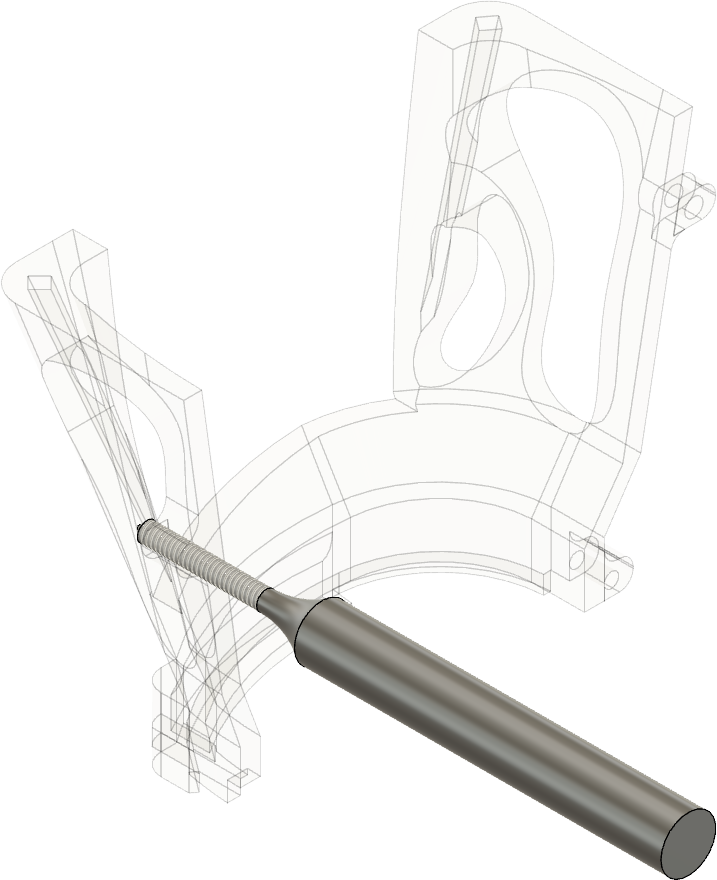

tapping hole

tapping hole

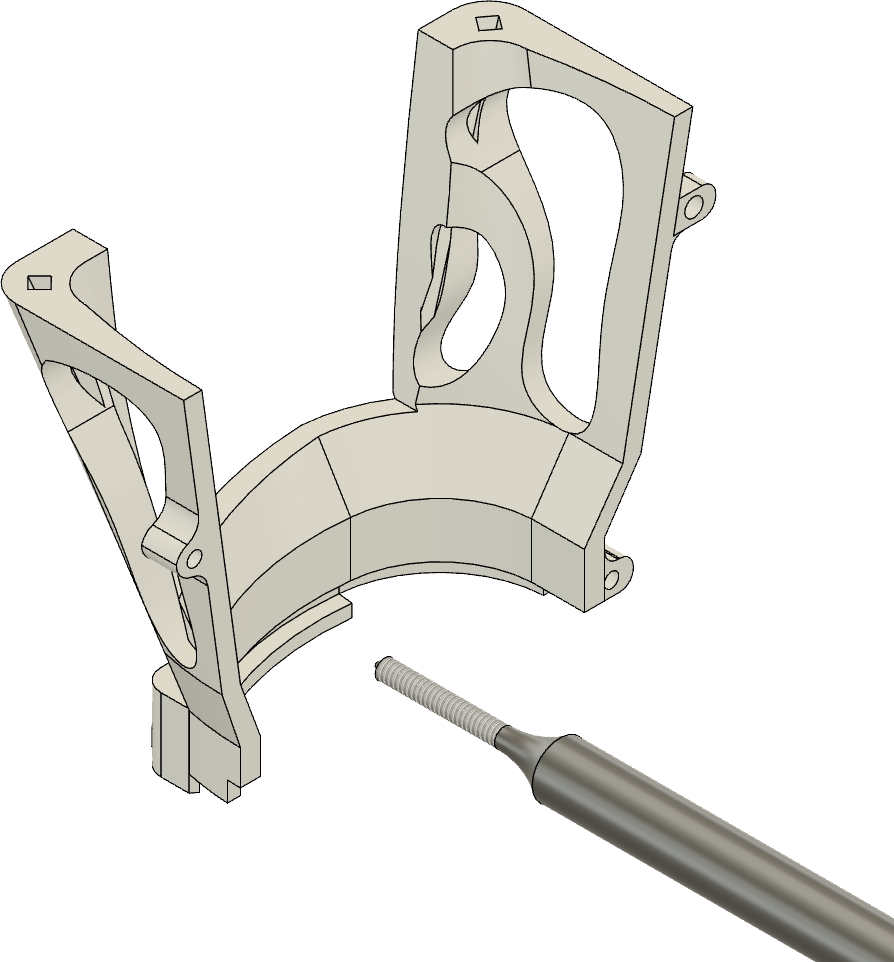

**1b - Tap the back holes (2x)**

right wall

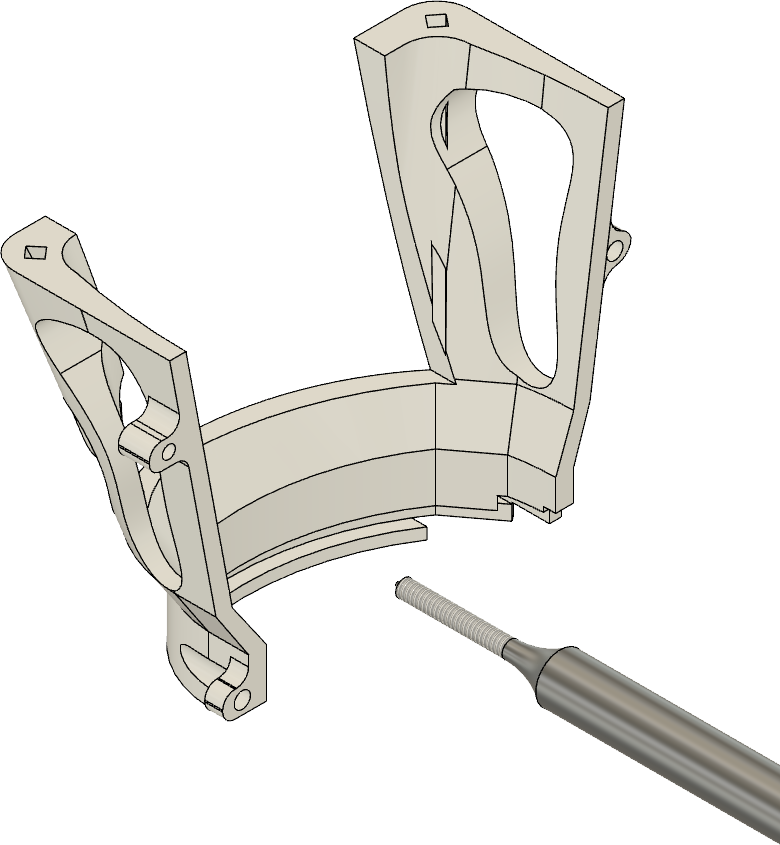

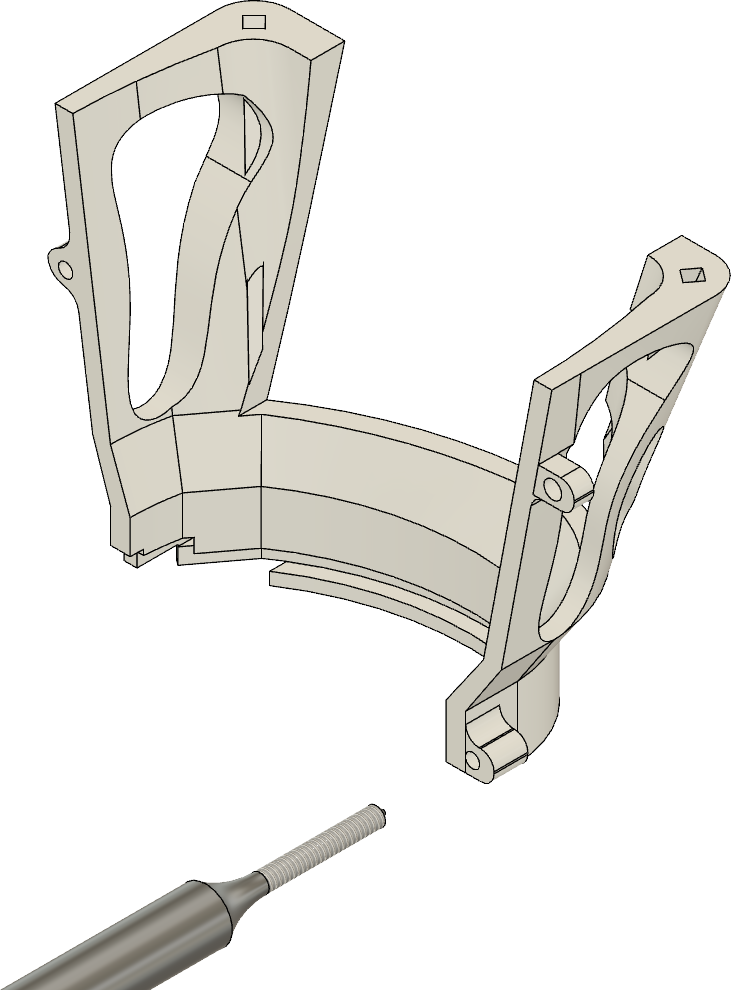

left wall

tapping hole

tapping hole

#### 2. Insert and glue metal bars into the walls.

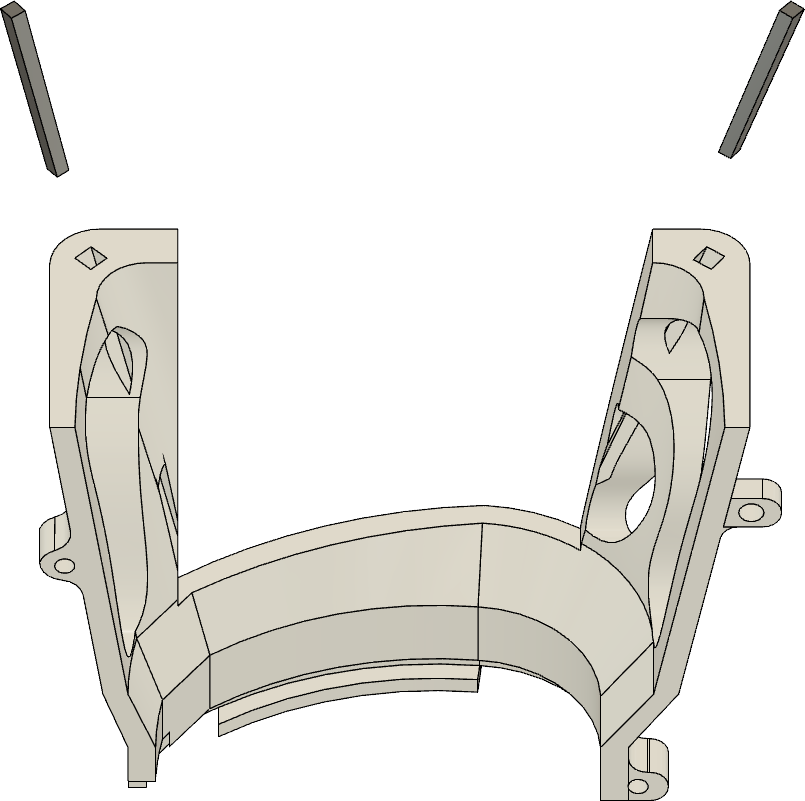

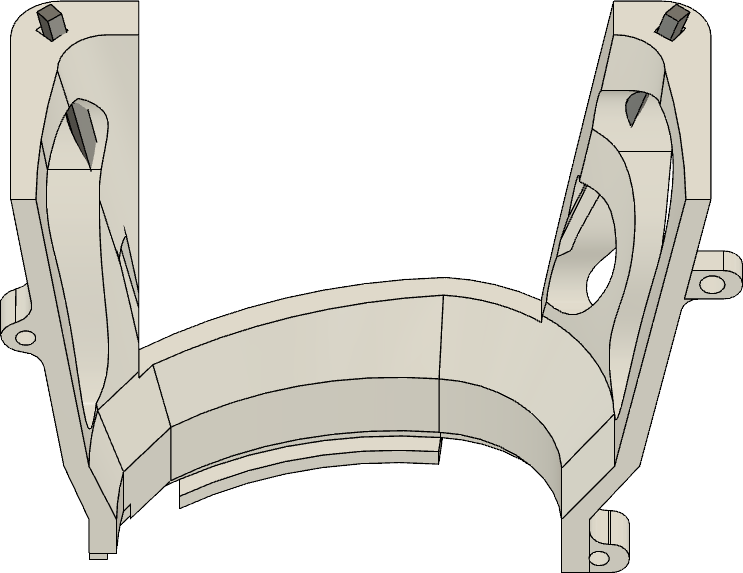

COMPLETED STEP

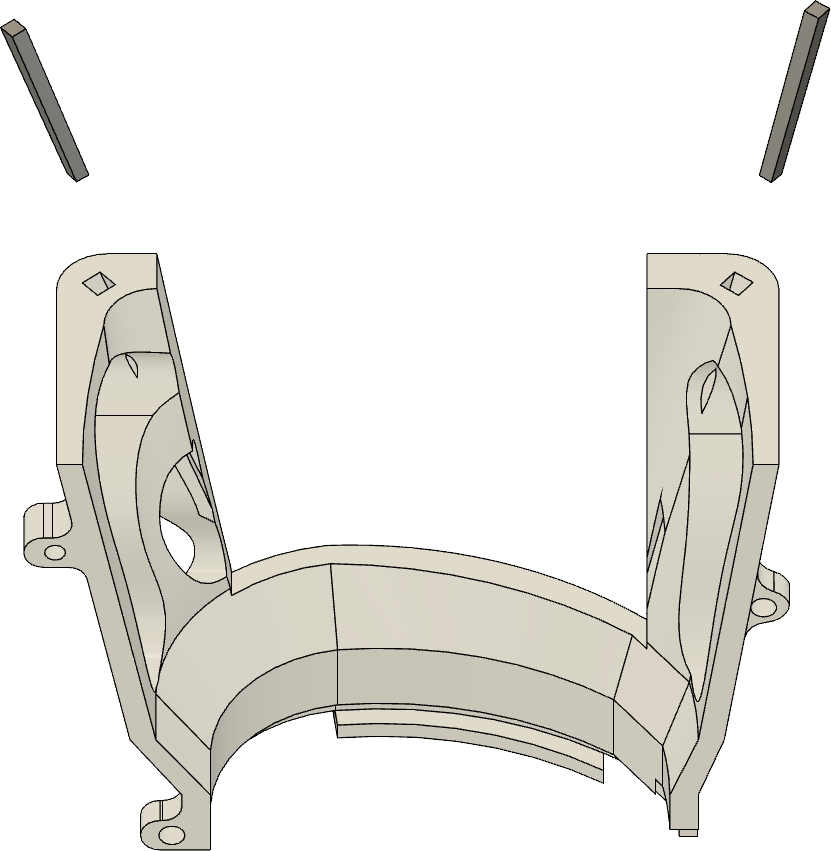

0.9 mm

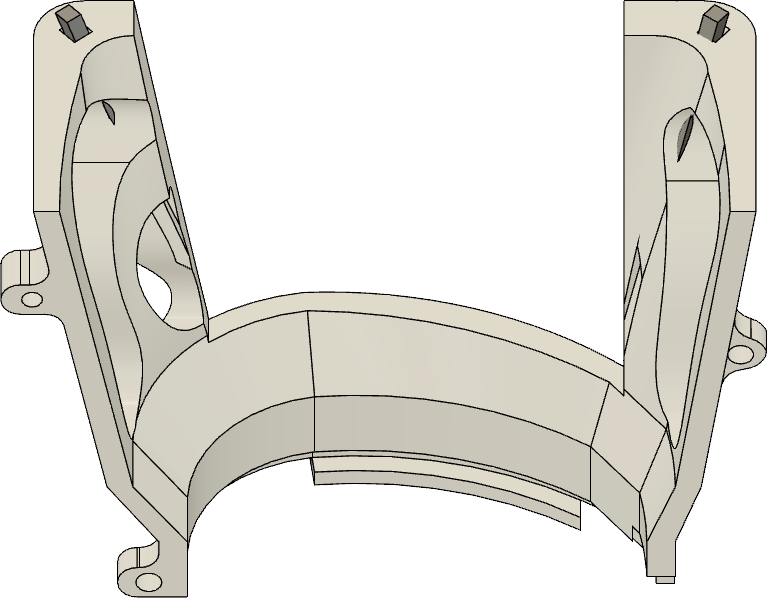

0.9 mm

0.7 mm0.7

mm

#### 3. Solder metal bar to the walls.

COMPLETED STEP

solder	solder

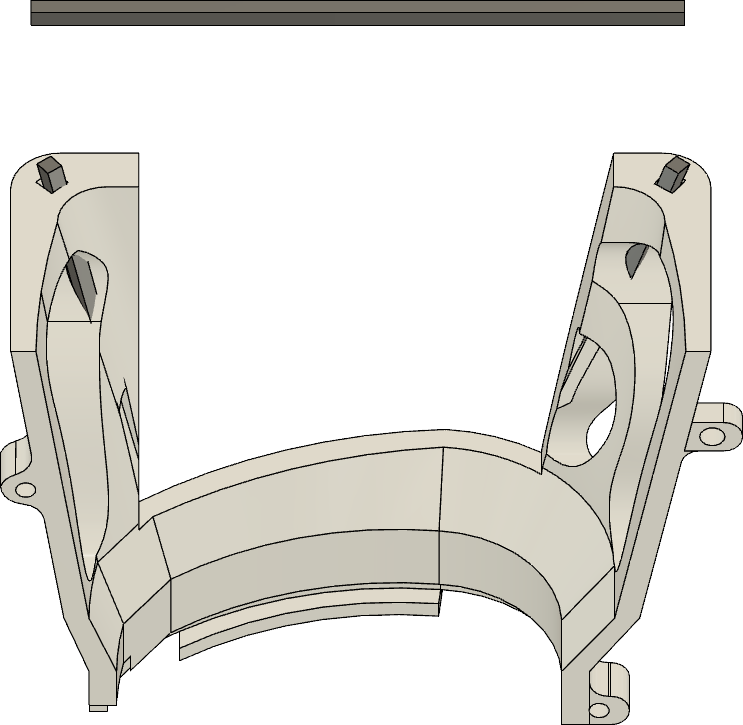

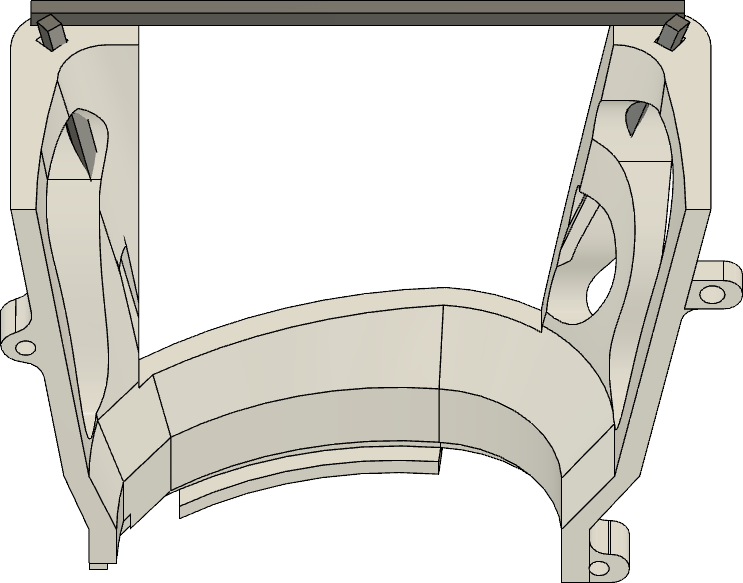

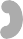

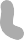

#### 4. Attach copper mesh to the wall (use solder and dental acrylic).

COMPLETED STEP

dental acrylic

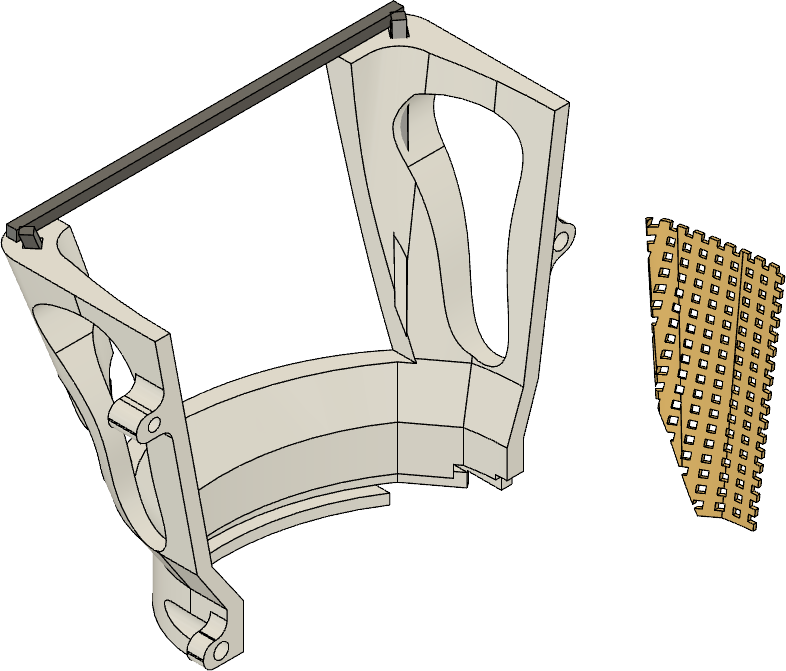

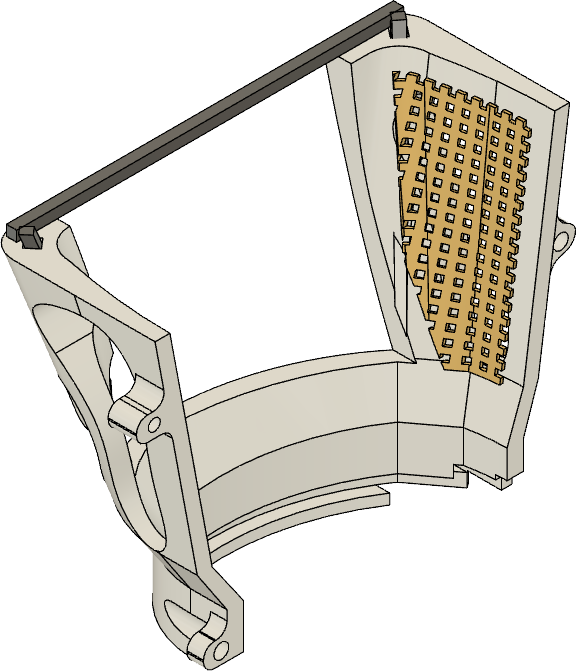

dental acrylic

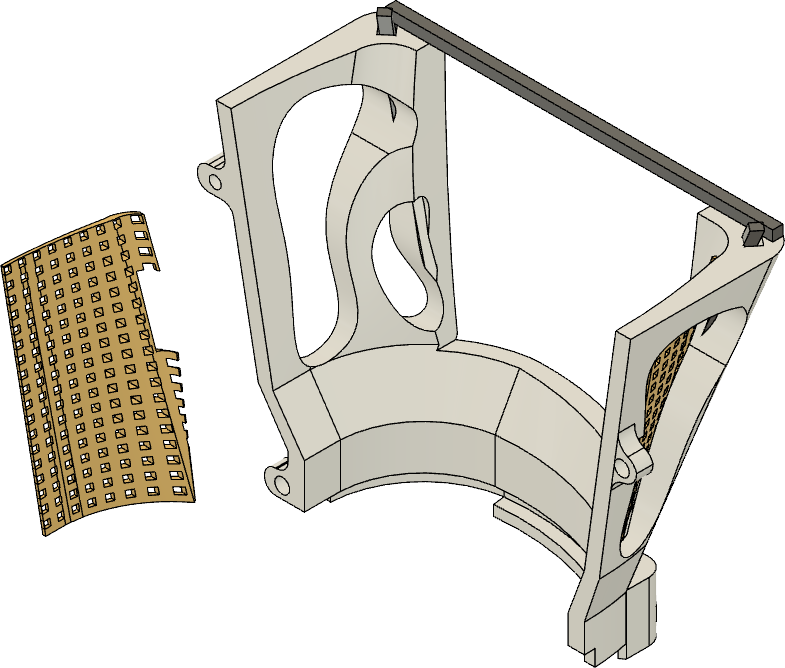

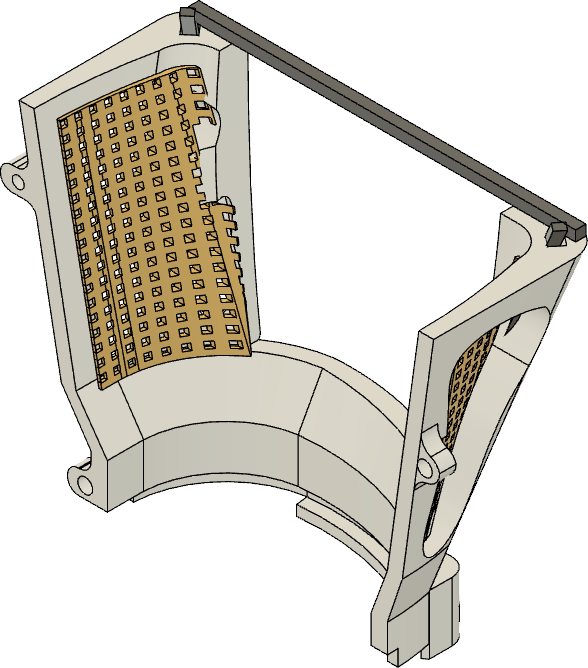

solder

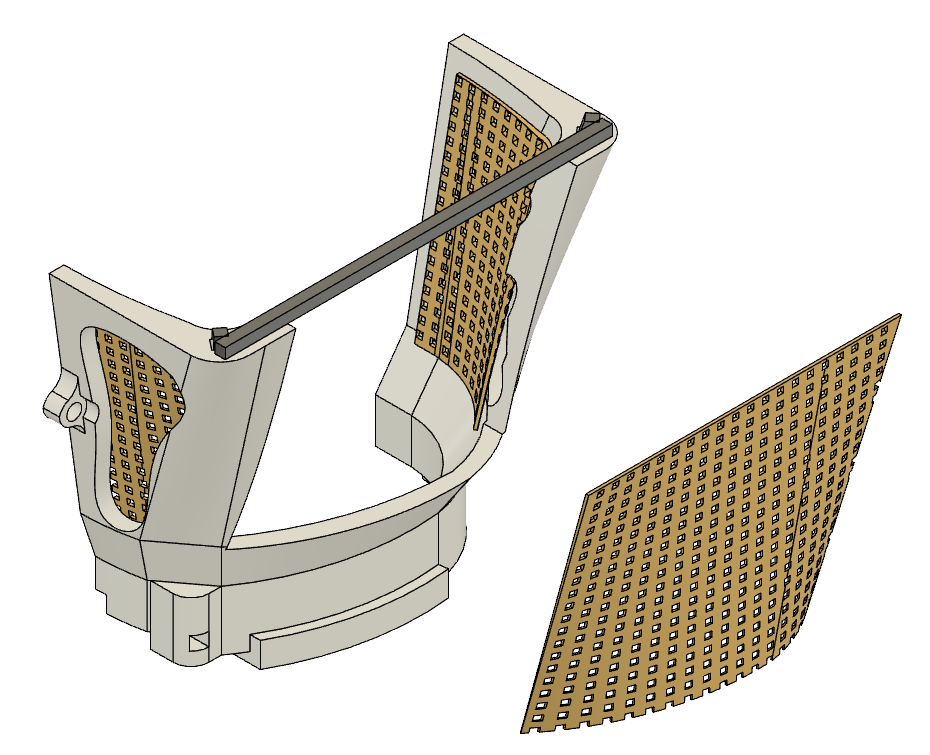

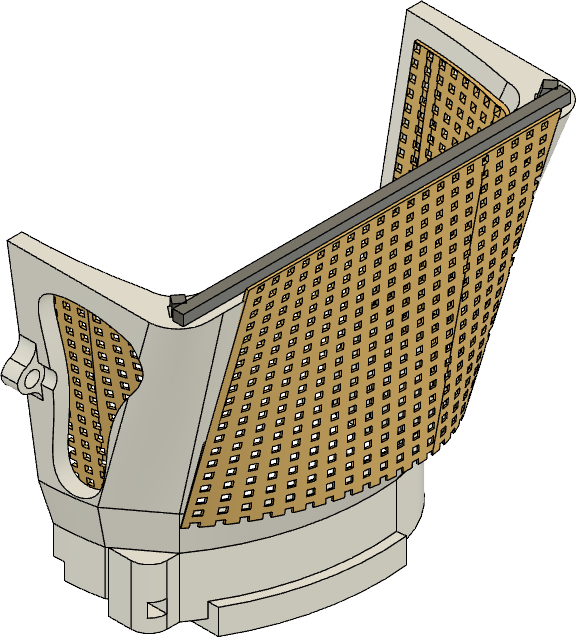

dental acrylic

pg. 3

#### 5. Attach walls to base.

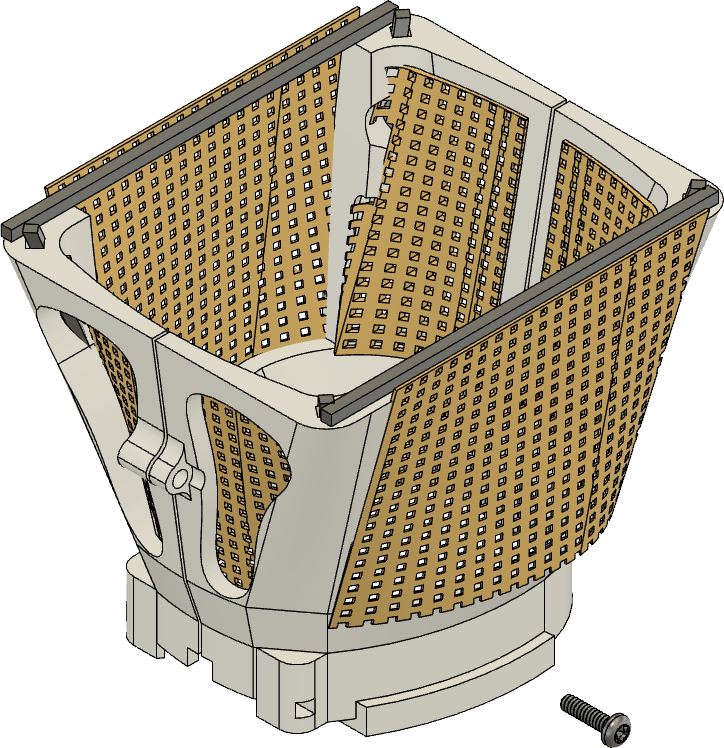

COMPLETED STEP

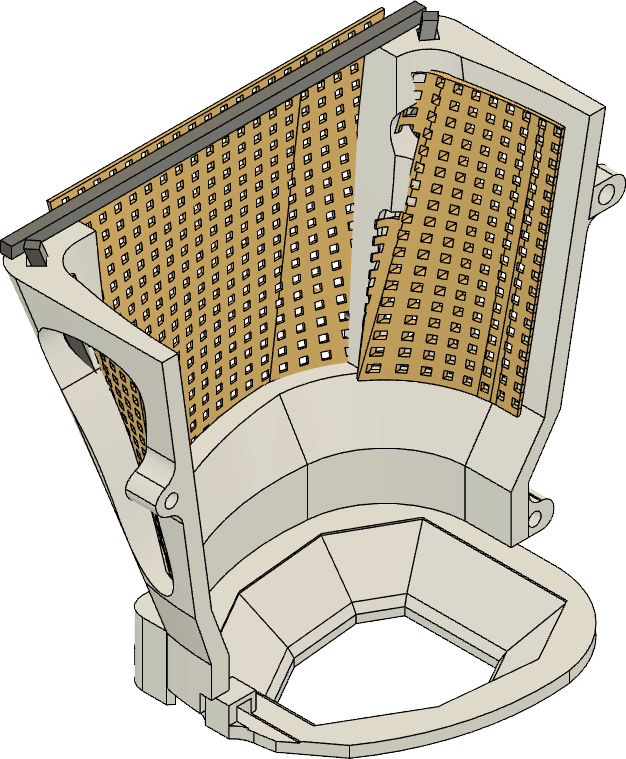

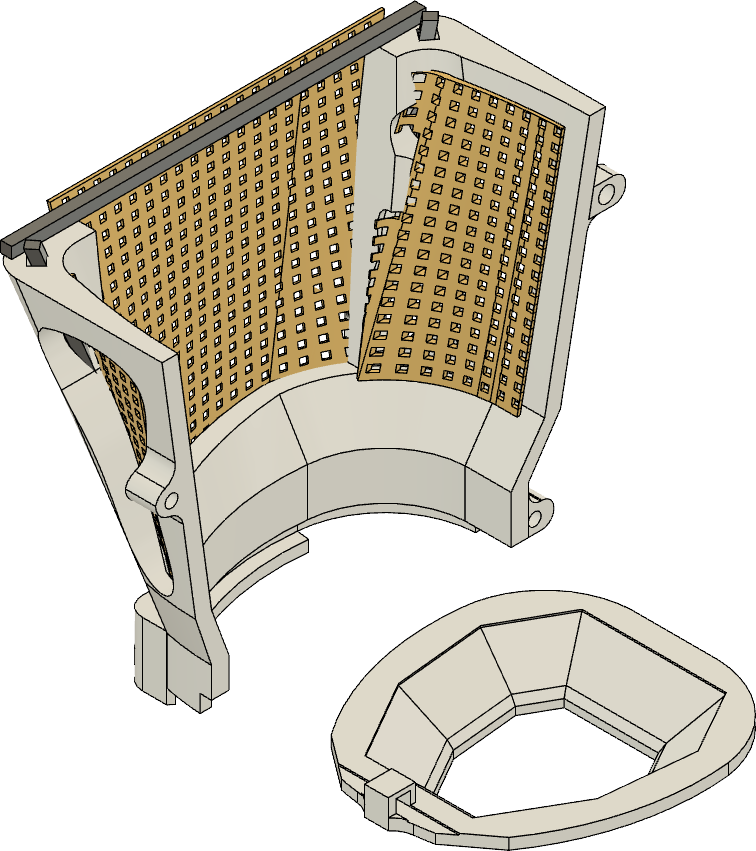

#### 6. Use metal bar during surgery for temporary fixation.

COMPLETED STEP

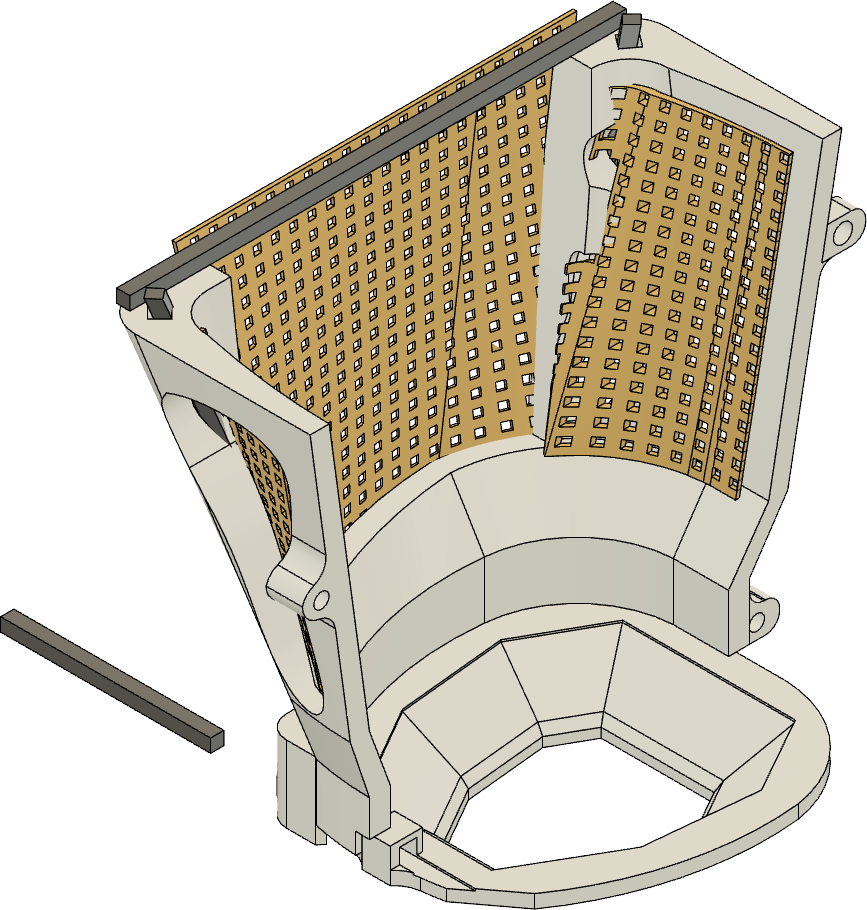

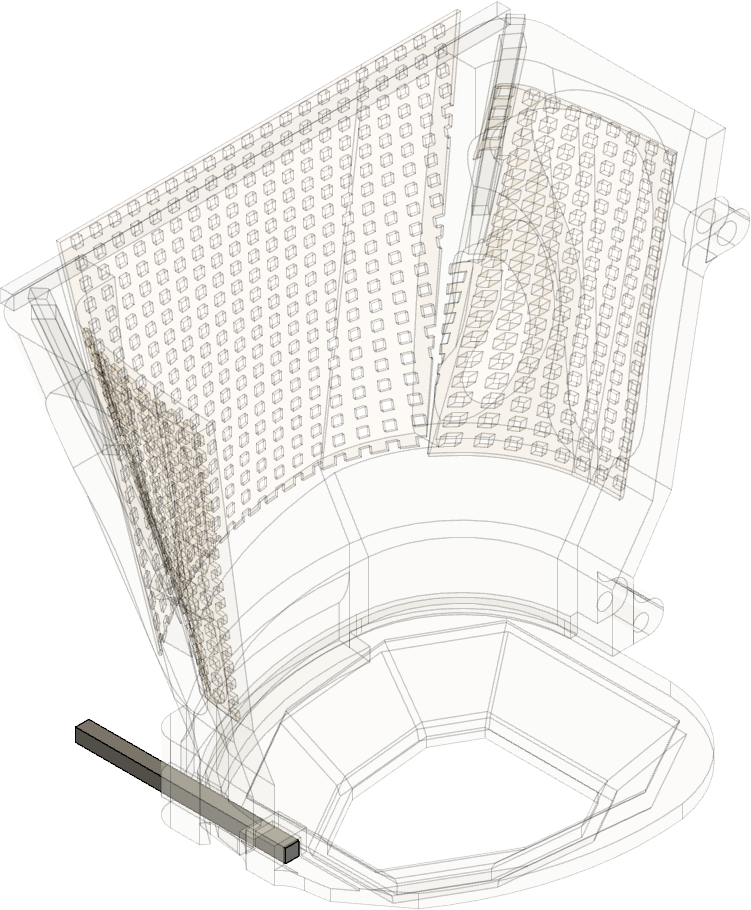

#### 7. Close walls using 00-120 screws in the front.

COMPLETED STEP

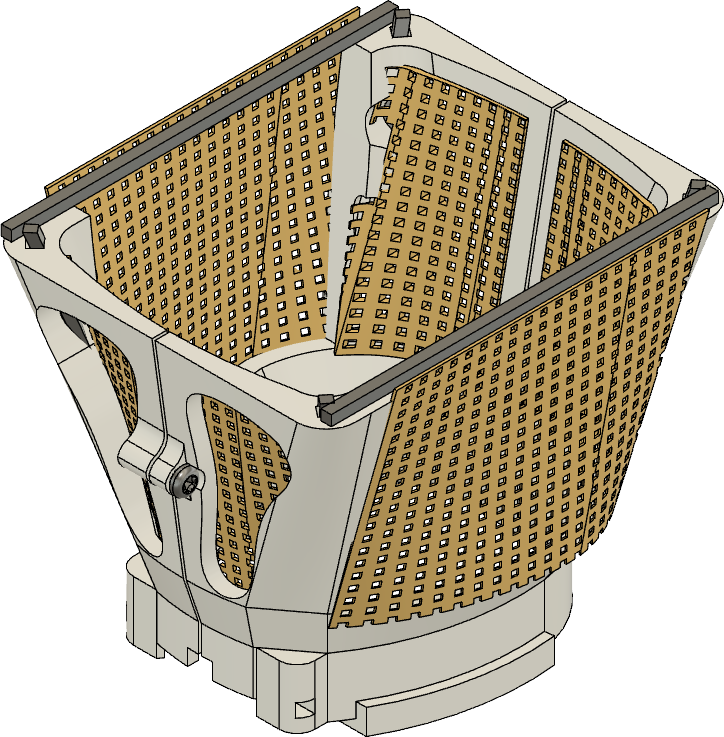

#### 8. Close walls using 00-120 screws in the back.

COMPLETED STEP

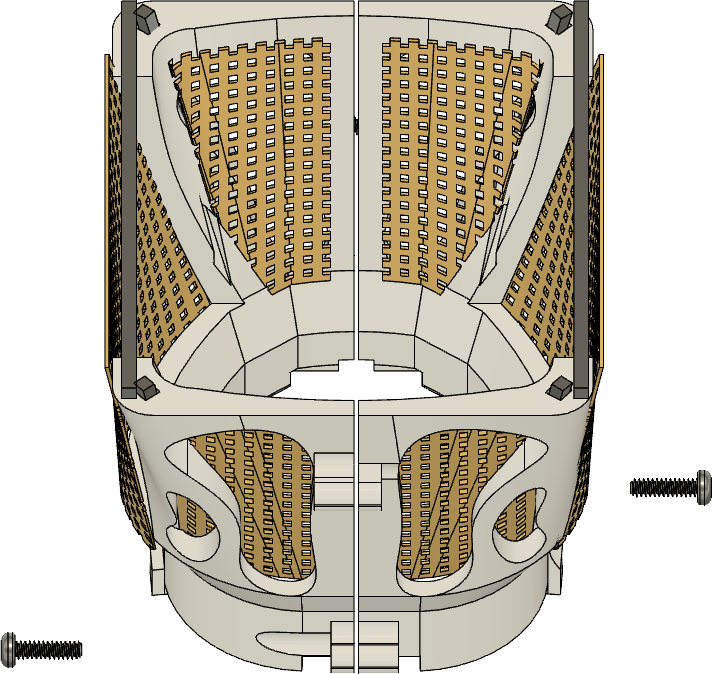

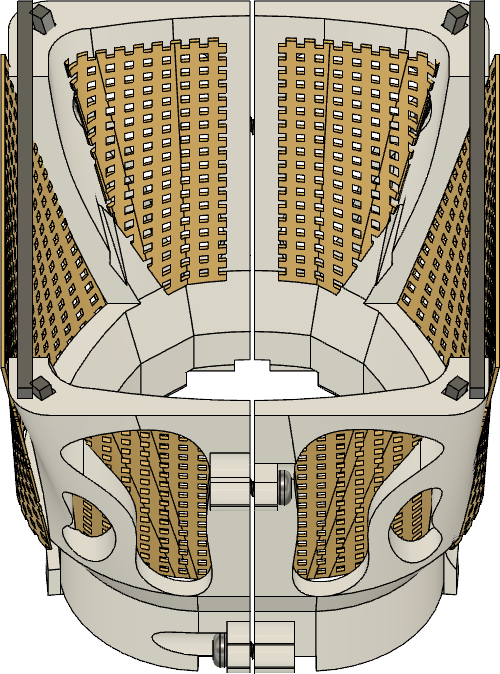

**Fully assembled mouse cap.**

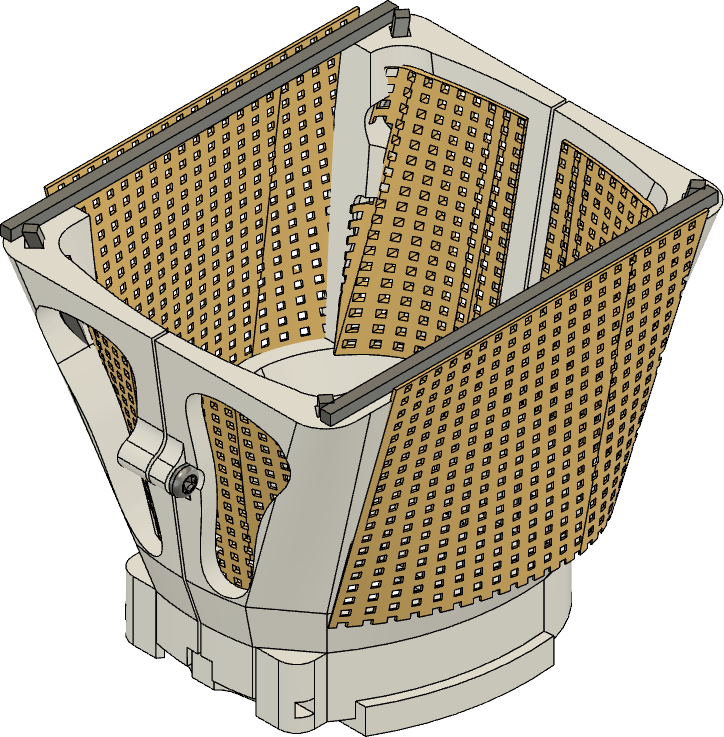

**3D-printed rat cap**

Design name: rat_hat_v8

More information is available at:

- https://github.com/misiVoroslakos/Rat_cap
- or contact me (Misi Voroslakos) directly at

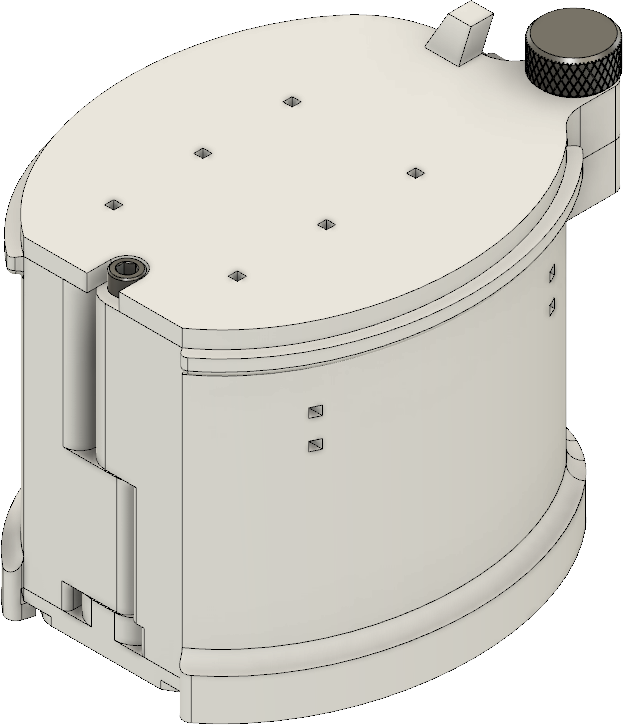

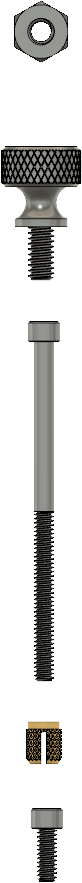
M2 nut (thin)

M2 thumb screw

00-80 screw, 1”

Wall

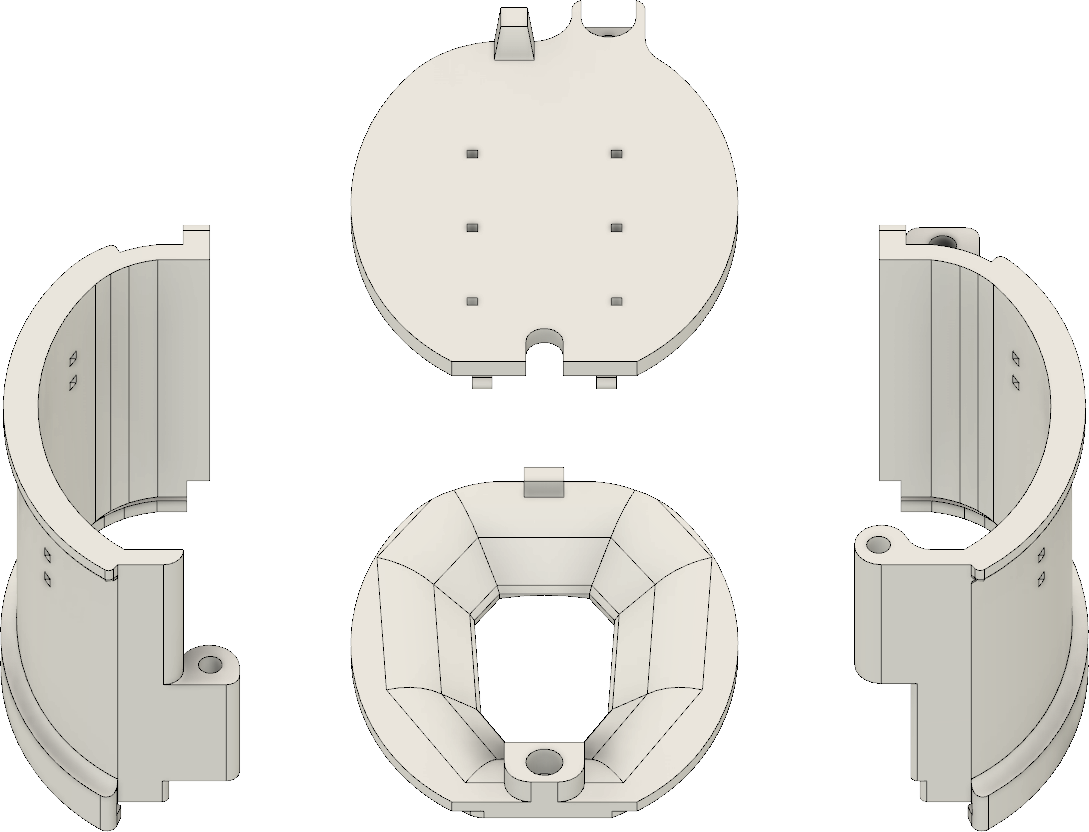

Lid

Base

Wall

00-80 insert

00-80 screw, 5/32” (2x)

#### 1. Glue 00-80 insert into base.

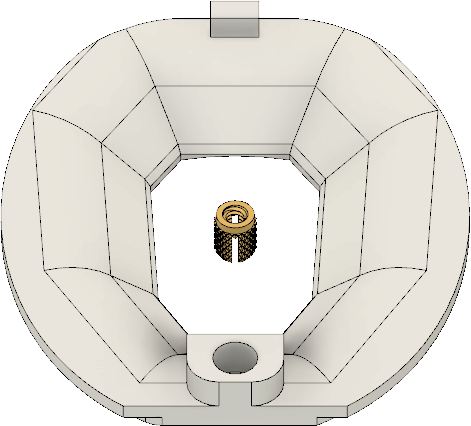

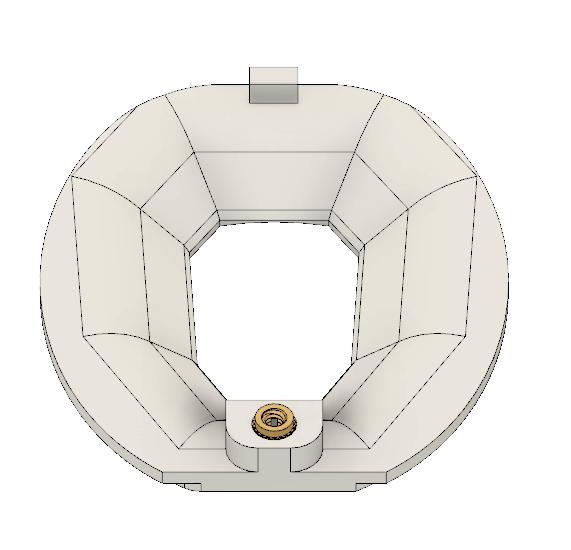

COMPLETED STEP

2. **Tap walls (00-80 tap).**

#### left wall

COMPLETED STEP

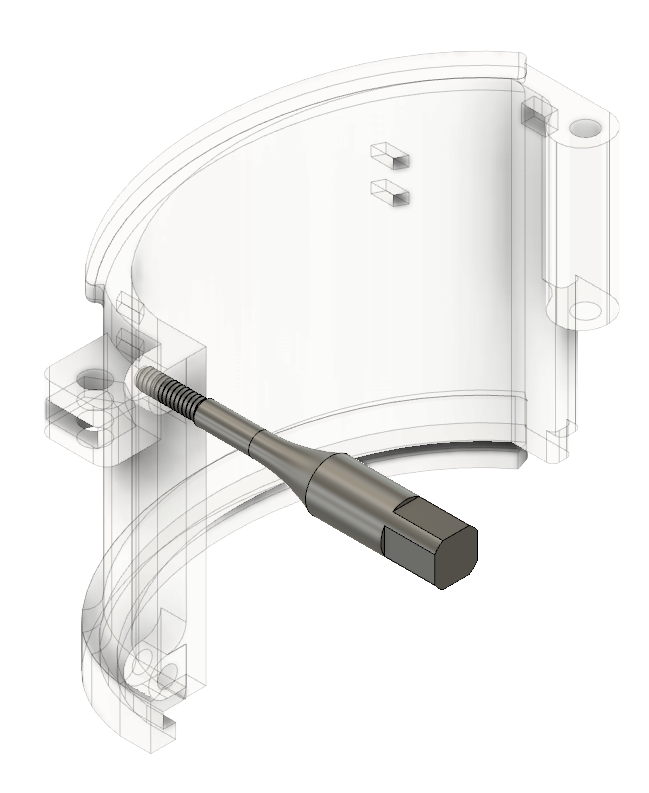

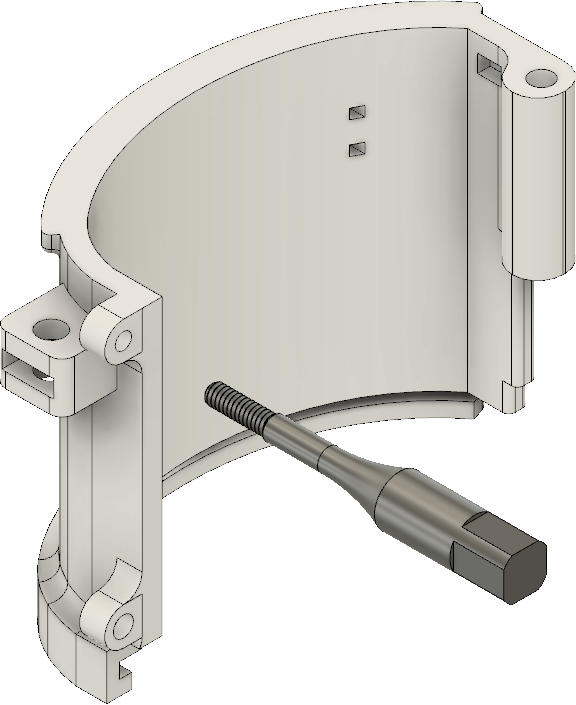

#### right wall

COMPLETED STEP

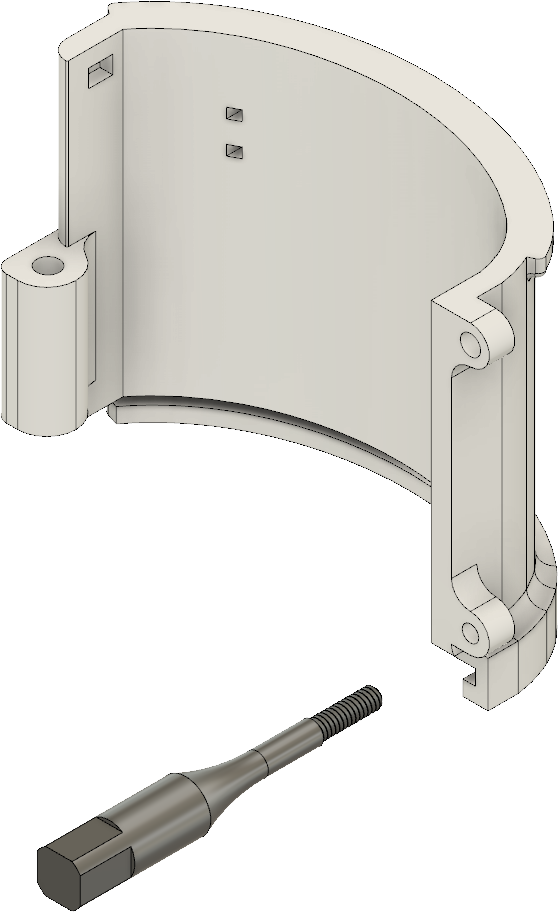

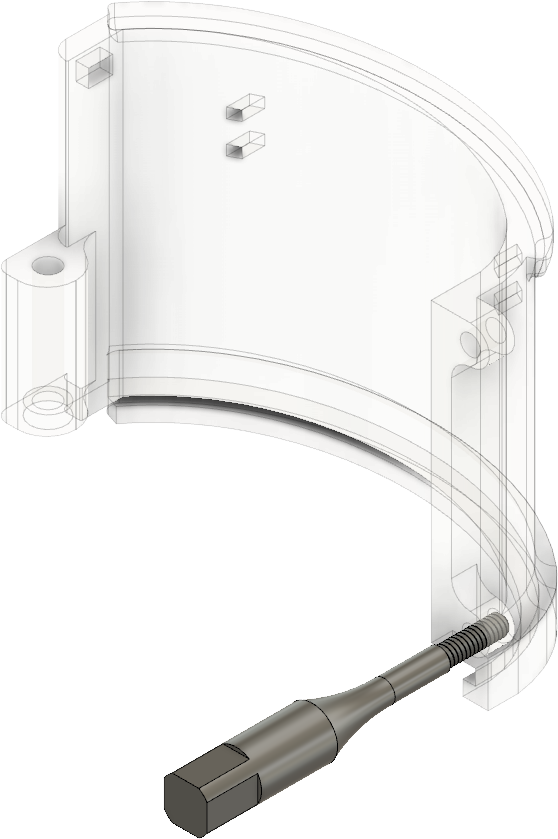

#### 3. Insert M2 nut in wall.

COMPLETED STEP

## M2

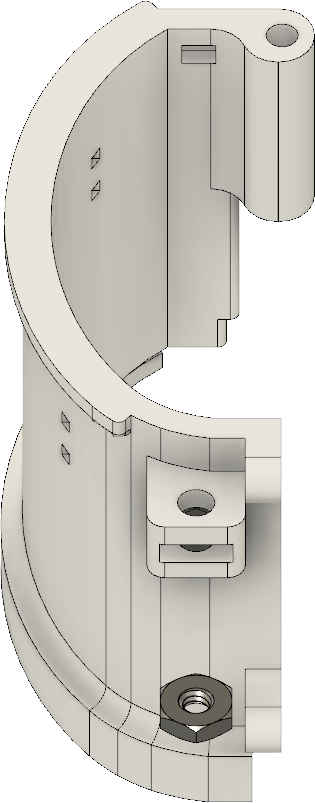

**nut**

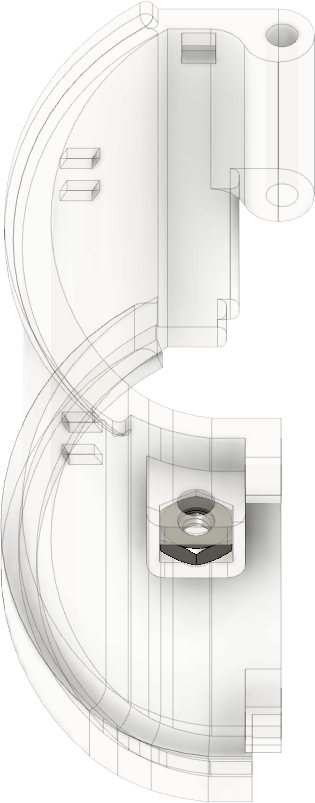

4. **Attach copper mesh.**

COMPLETED STEP

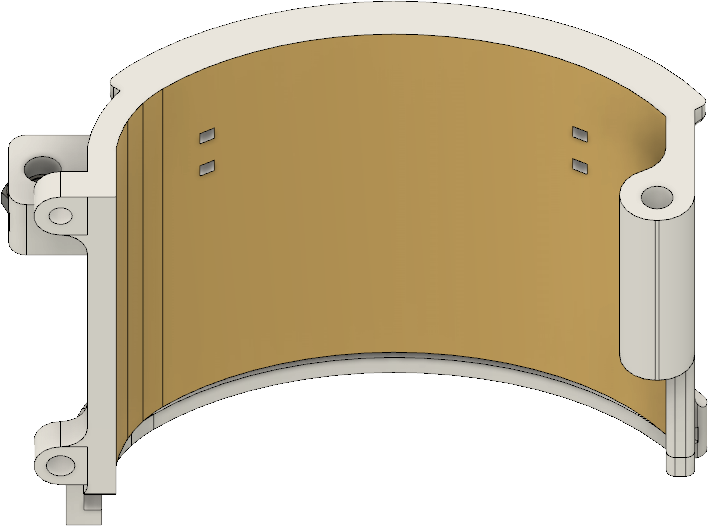

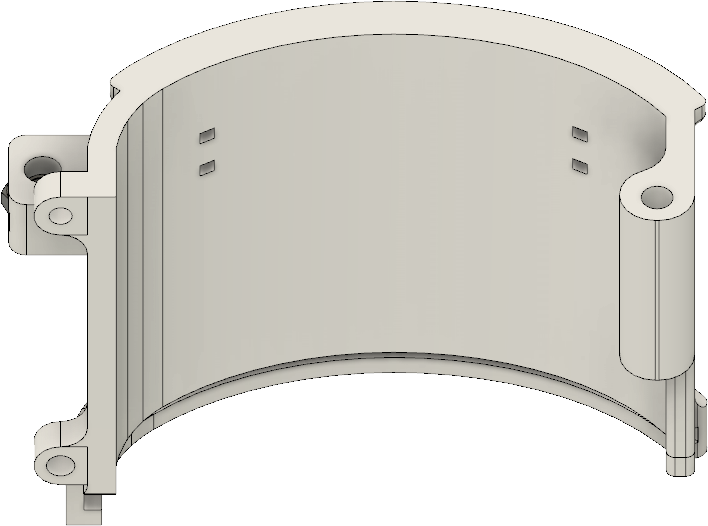

copper tape

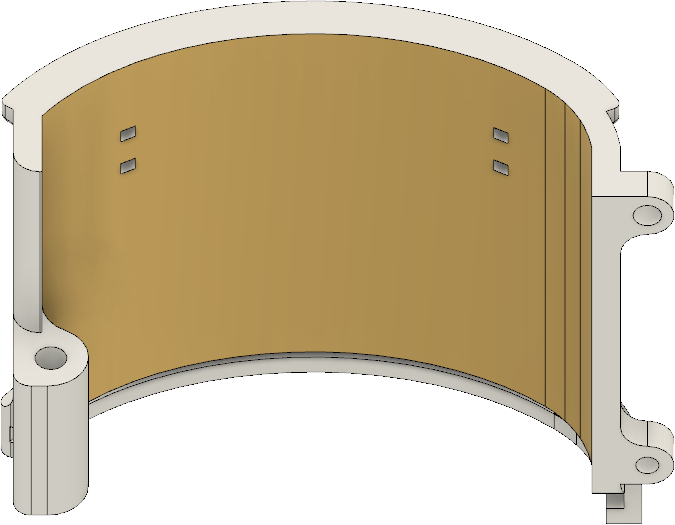

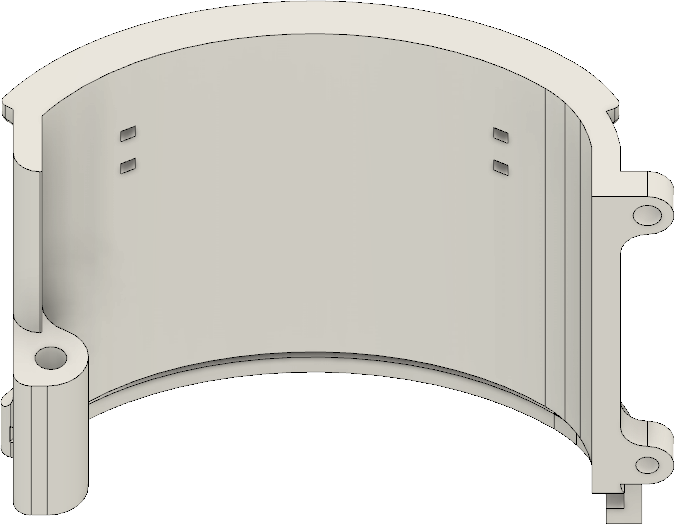

copper tape

#### 5. Insert and glue metal bars to walls.

COMPLETED STEP

metal bars

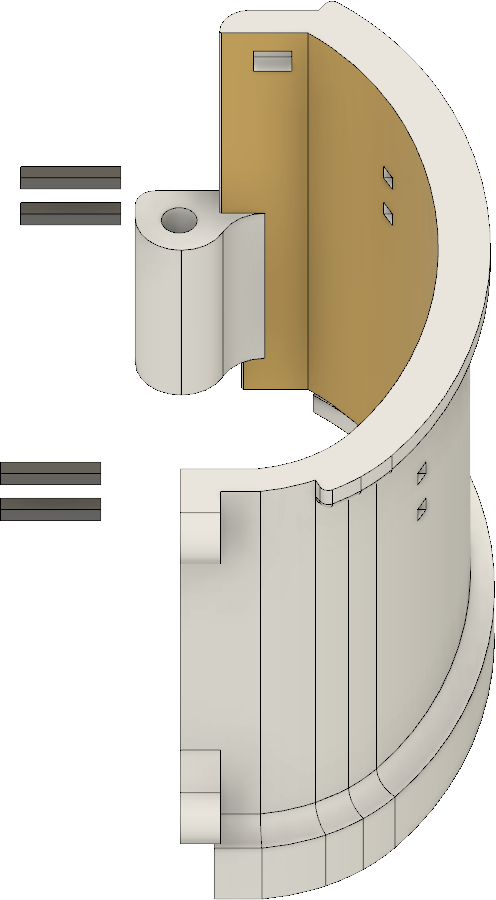

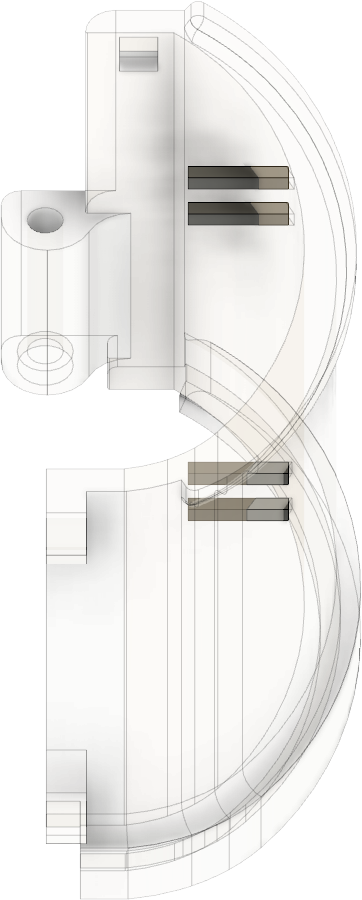

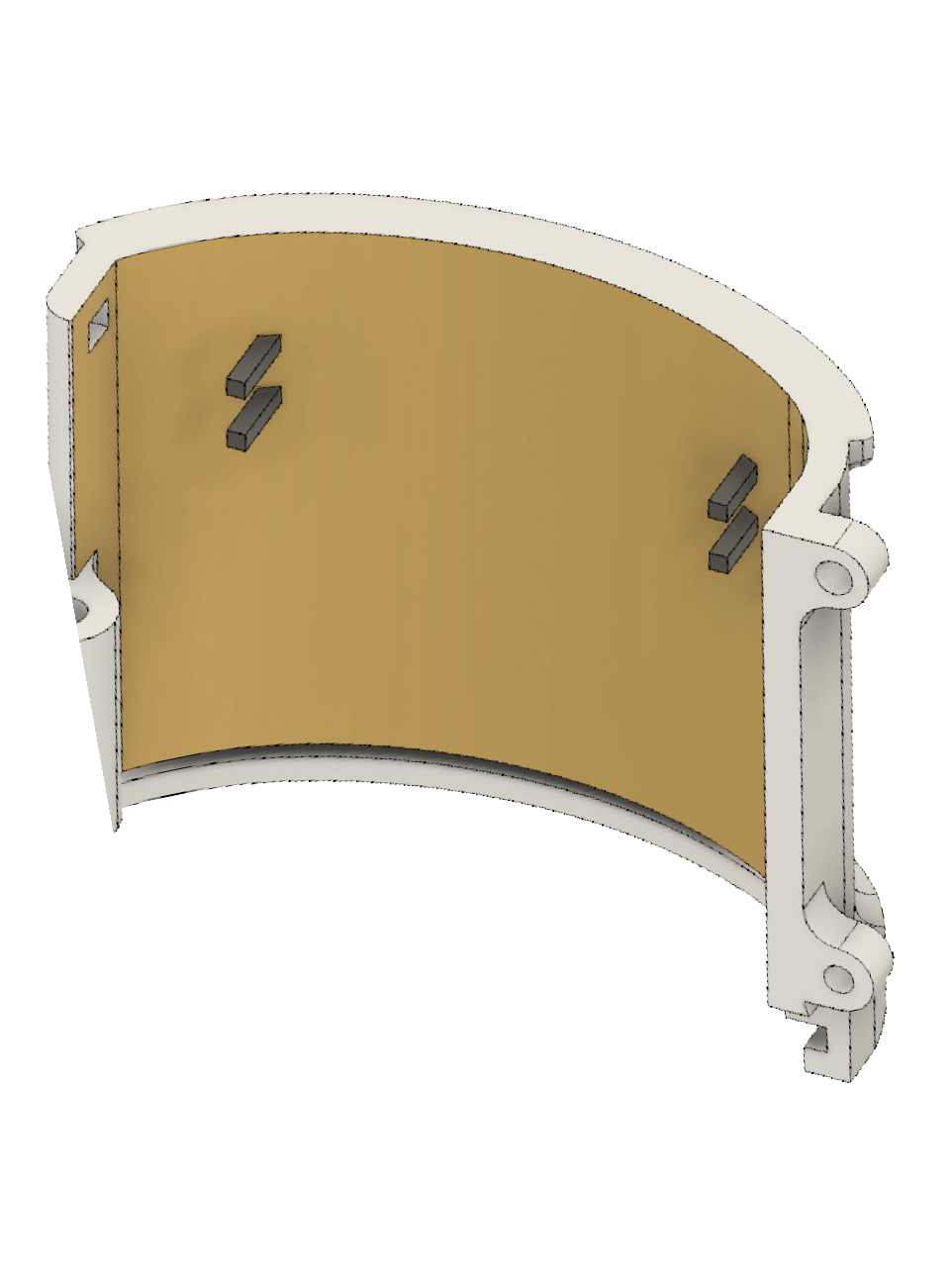

COMPLETED STEP

metal

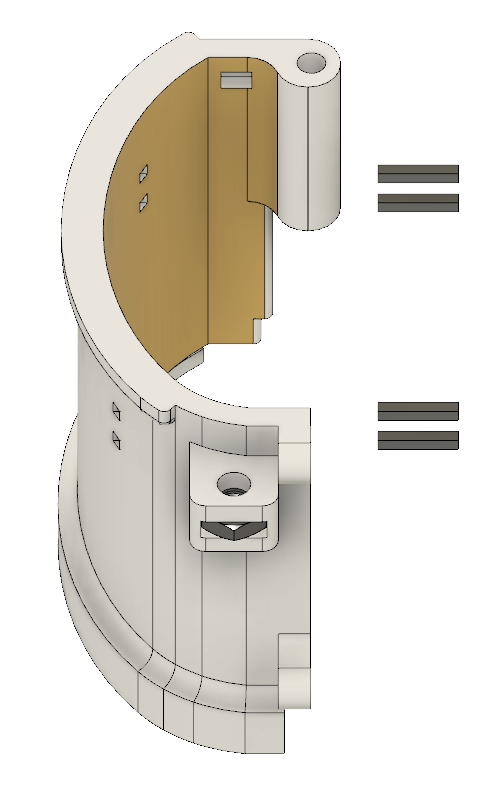

bars

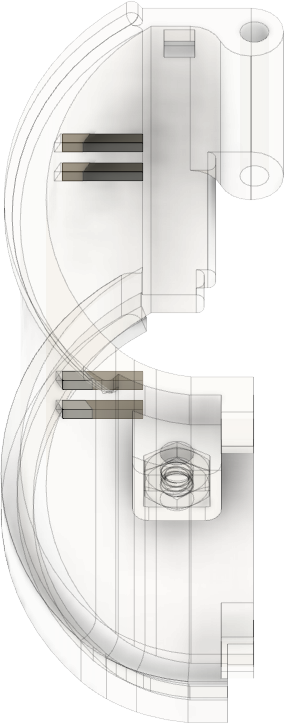

These steps are also available in video format (check the github page).

#### 6. Attach walls to base.

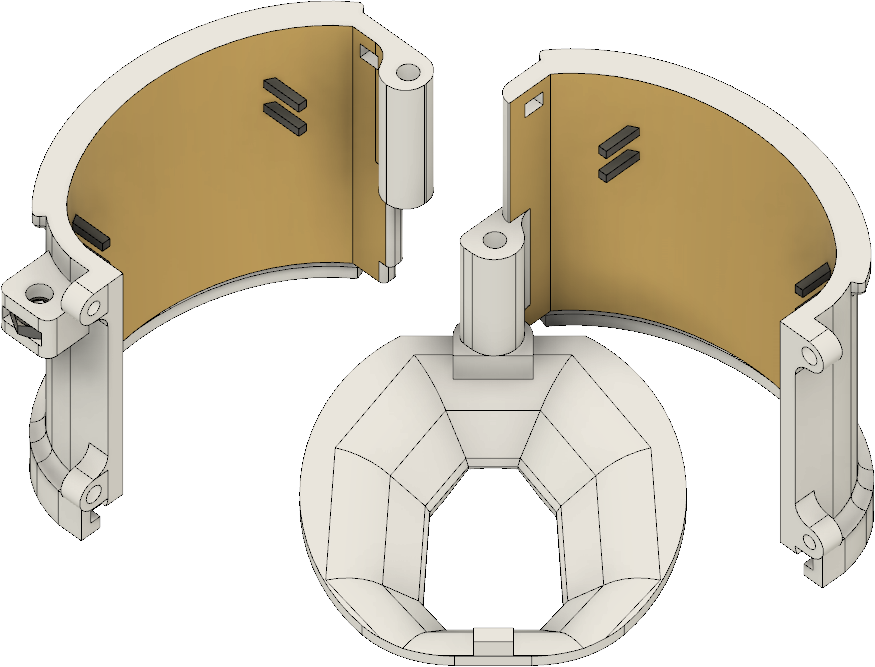

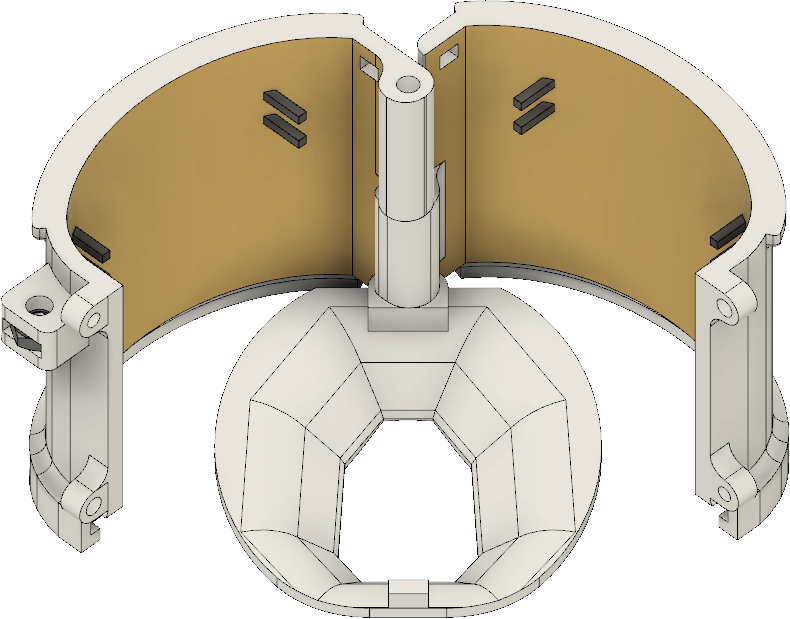
COMPLETED STEP

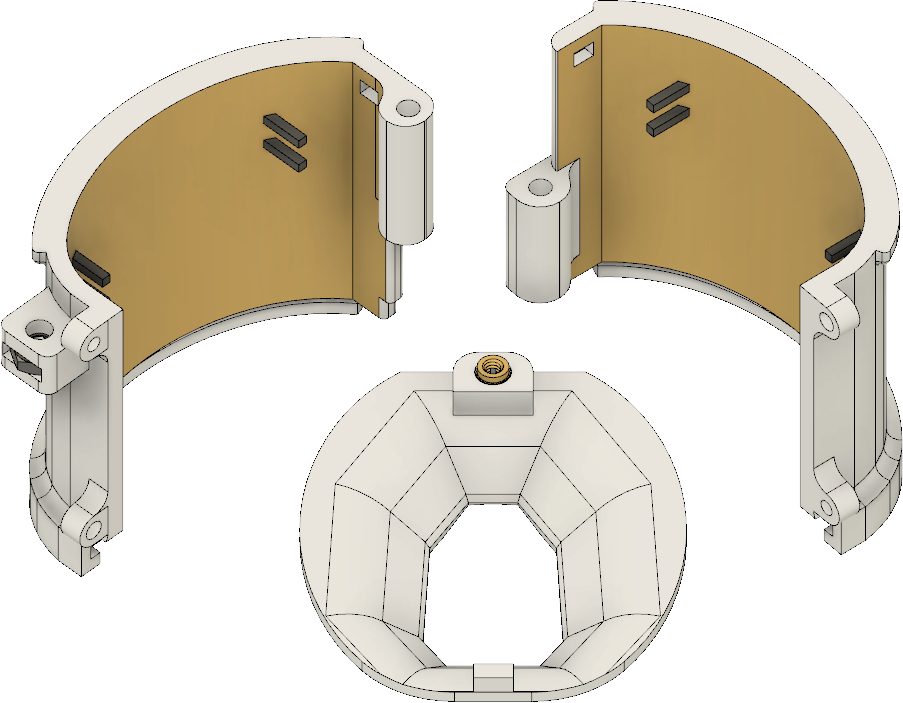

#### 7. Close the walls and insert 00-80 screw (1”).

COMPLETED STEP

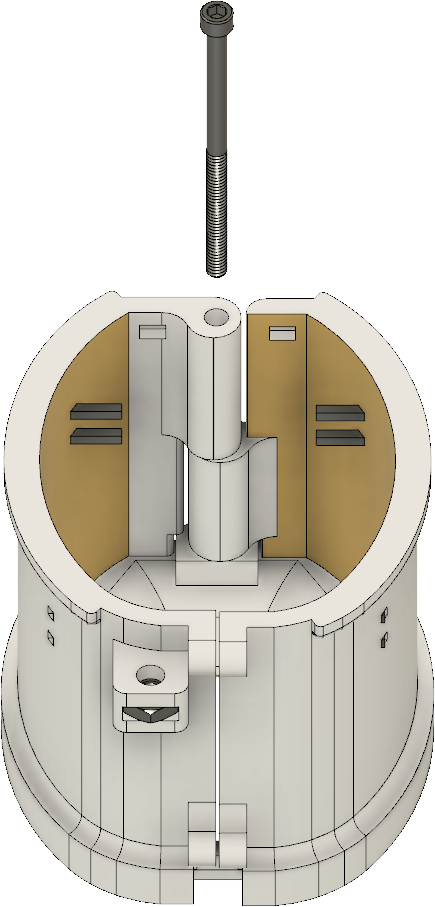

00-80

screw (1”)	00-80 screw (1”)

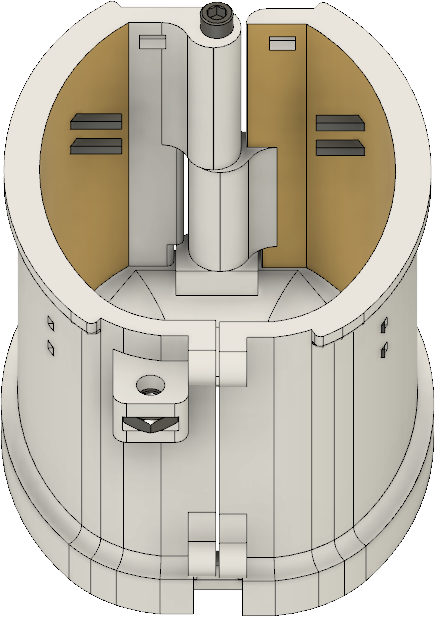

#### 8. Insert 00-80 screw (5/32”).

COMPLETED STEP

00-80 screw (5/32”)

00-80 screw

(5/32”)	00-80 screw

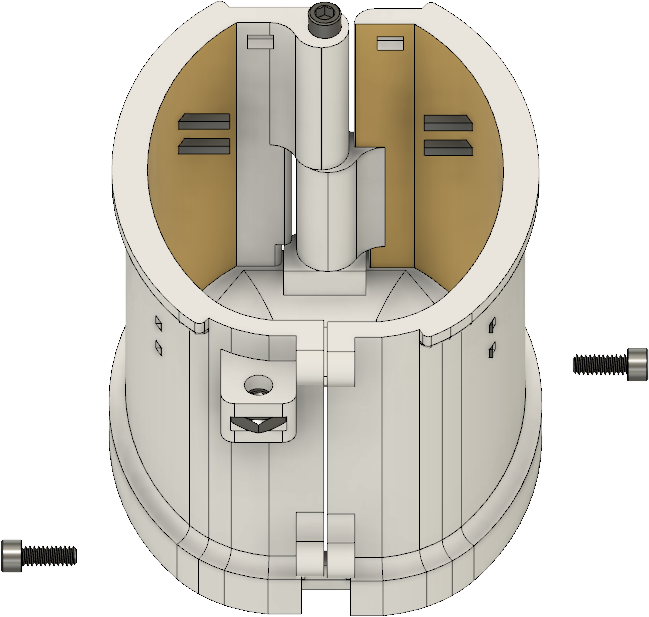

(5/32”)

pg. 5

#### 9. Insert thumb screw (M2).

COMPLETED STEP

M2 thumb screw

M2 thumb screw

#### Fully assembled design

This edge helps to keep coband attached to the cap in case you don’t want to use the plastic lid.

pg. 6
